# Chronic exposure to glucocorticoids amplifies inhibitory neuron cell fate during human neurodevelopment in organoids

**DOI:** 10.1101/2024.01.21.576532

**Authors:** Leander Dony, Anthi C. Krontira, Lea Kaspar, Ruhel Ahmad, Ilknur Safak Demirel, Malgorzata Grochowicz, Tim Schaefer, Fatema Begum, Vincenza Sportelli, Catarina Raimundo, Maik Koedel, Marta Labeur, Silvia Cappello, Fabian J. Theis, Cristiana Cruceanu, Elisabeth B. Binder

## Abstract

Disruptions in the tightly regulated process of human brain development have been linked to increased risk for brain and mental illnesses. While the genetic contribution to these diseases is well established, important environmental factors have been less studied at molecular and cellular levels. In this study, we used single-cell and cell-type-specific techniques to investigate the effect of glucocorticoid (GC) exposure, a mediator of antenatal environmental risk, on gene regulation and lineage specification in unguided human neural organoids. We characterized the transcriptional response to chronic GC exposure during neural differentiation and studied the underlying gene regulatory networks by integrating single-cell transcriptomics-with chromatin accessibility data. We found lasting cell type-specific changes that included autism risk genes and several transcription factors associated with neurodevelopment. Chronic GCs influenced lineage specification primarily by priming the inhibitory neuron lineage through key transcription factors like PBX3. We provide evidence for convergence of genetic and environmental risk factors through a common mechanism of altering lineage specification.

## Introduction

Human neurodevelopment is a tightly regulated process starting early in embryogenesis that choreographs cellular proliferation, migration, differentiation, and synaptogenesis. The precise timing and sequence of these events are essential to establish neural circuits that govern cognitive, emotional, and behavioral functions. Deviations from this program have been linked to a spectrum of neurodevelopmental and psychiatric disorders, including autism spectrum disorders (ASD) and schizophrenia. This is supported by the strong enrichment of genes carrying genetic variants associated with these disorders in molecular and cellular pathways essential for neurodevelopment ^1,2,3^. Modeling the effects of associated deleterious variants in transgenic animals and induced pluripotent stem cell-derived model systems supports their impact on neurodevelopment ^4,5^.

While these disorders have a large genetic component, with heritability estimates from twin studies around 75 % ^6^, environmental risk factors acting during pregnancy such as chemicals ^7^, infections ^8^, perinatal complications ^9,10^, and exposure to glucocorticoids (GCs) ^11^ have also been implicated in increasing disease risk by impacting neurodevelopment. With this work, we aimed to elucidate the contribution of one such prenatal environmental factor, GCs, which are steroid hormones with critical endogenous roles in normal brain development and important pharmacological applications during pregnancy ^11^. Activation of the GC receptor in the developing brain plays an essential role in neurogenesis, neuronal migration, synaptogenesis, and modulation of neuronal plasticity ^11^. Perturbations in GC signaling during critical periods of brain development have been proposed to lead to long-lasting alterations in brain structure and function, potentially contributing to the pathogenesis of psychiatric disorders ^12,13,14^. Animal models and human studies have provided compelling evidence for adverse effects of GC excess during gestation on cognitive and emotional development. During pregnancies at risk of premature labor, synthetic GCs such as betamethasone and dexamethasone are routinely used to promote lung maturation in the unborn child. Large epidemiological studies have linked such antenatal exposure to synthetic GCs to altered risk for childhood mental and behavioral problems ^15,16,17^. An increase in risk is observed mainly when synthetic GCs are administered later in pregnancy, while the opposite was observed for extremely preterm babies born at less than 28 weeks gestation ^18,19^. Given that more than ten percent of babies (13 million) were born prematurely worldwide in 2020, this is a sizable environmental factor for brain development ^20^.

Despite the strong evidence for harmful effects from epidemiological studies, there is limited evidence on the underlying molecular and cellular mechanisms ^11,21^. Using human neural organoids as models of early brain development, we could recently show that GCs elicit cell-type specific transcriptional responses and that there is a highly significant enrichment of genes associated with neurodevelopmental delay, ASD, and more common psychiatric disorders among those regulated by GCs, especially in neurons ^22^. This suggests a convergence of the molecular pathways underpinning genetic and environmental risk factors for neuropsychiatric disorders.

The advent of human neural organoids now allows us to better link risk genes for neurodevelopmental disorders to molecular and cellular mechanisms. Data from cerebral organoids derived from induced pluripotent stem cells from patients with idiopathic ASD or carrying rare coding or copy number variants associated with ASD suggest that cell fate specification is altered by risk genes carrying deleterious mutations, with particular support for alterations in the proportions of excitatory vs. inhibitory neuronal lineages or dorsal vs ventral arealization ^23,24,25,26,27^. These findings are further supported by results from a CRISPR-human organoids-single-cell RNA sequencing (CHOOSE) system, in which 36 high-risk autism spectrum disorder genes were perturbed. These risk genes were selected among other autism risk genes for their relation to transcriptional regulation, and their perturbation uncovered consistent effects on cell fate determination, mainly lowering the dorsal-to-ventral ratio at different levels ^28^.

With this manuscript, we aim to address whether the convergence of risk genes for neuropsychiatric and neurodevelopmental disorders and genes regulated by antenatal exposure to GC, an environmental risk factor for childhood mental and neurodevelopmental disorders, extends beyond the molecular context to alteration of cell fate specification. For this, we exposed unguided and regionalized neural organoids to a chronic administration of the synthetic GC dexamethasone, as it is used clinically in pregnancies at risk of premature labor. We assessed cell type-specific responses with single-cell RNA sequencing (scRNA-seq) and single-cell Assay for Transposase-Accessible Chromatin using sequencing (scATAC-seq) analyses directly after the chronic administration and following a wash-out period. Our work extends beyond the existing literature by demonstrating a convergent impact of environmental and genetic risk factors for mental and neurodevelopmental disorders on cell fate determination in the developing brain.

## Results

### Chronic glucocorticoid exposure in neural organoids does not induce significant metabolic stress in cells

To elucidate the effects of GCs on neurodevelopmental processes, we designed an exposure paradigm in unguided neural organoids. We collected 70-day-old organoids, continuously exposed to GCs for ten days (Chr condition) and the matching vehicle control organoids (Veh condition), allowing us to measure the immediate transcriptional effects of chronic GC exposure. Furthermore, we were interested in understanding which transcriptional changes lasted for an extended period in organoids. For this, we collected organoids from the Veh and Chr conditions after an additional 20 days under standard culture conditions at day 90 (Veh-Veh and Chr-Veh conditions, respectively). Lastly, we wanted to understand the differences between the transcriptional effects of chronic and acute exposure. Therefore, we obtained samples at day 90, exposed to an additional 12-hour acute GC treatment before collection (Veh-Acu and Chr-Acu conditions). We duplicated all analyses using organoids derived from two human induced pluripotent stem cell (hiPSC) lines, Line 409b2 and Line FOK4, with four replicates in each of the six experimental conditions (Fig. 1).

**Fig. 1.**
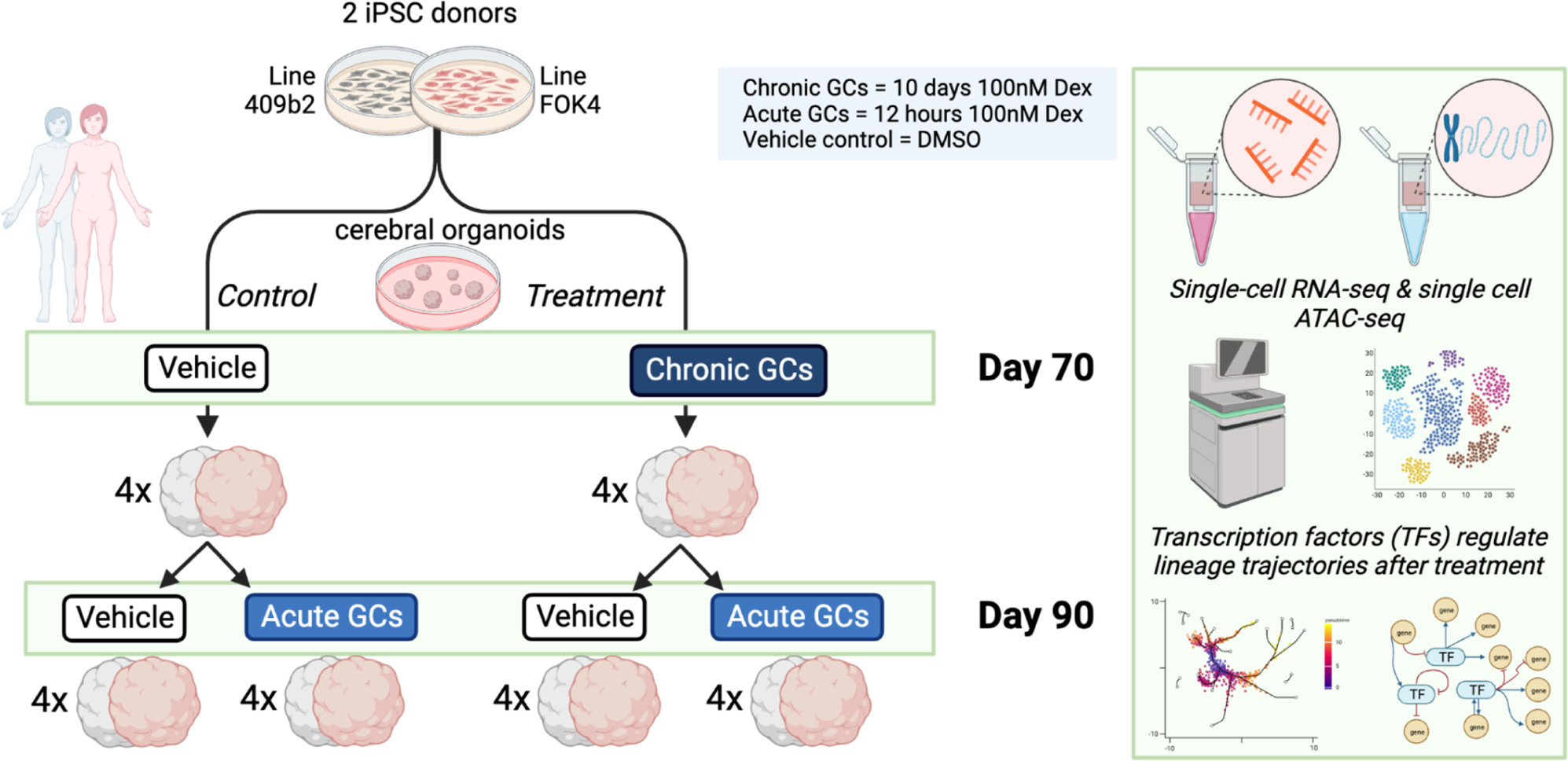
Overview of experimental setup and design. We designed six treatment conditions duplicated in organoids derived from two cell lines. We replicated each of these 12 conditions using four samples. We collected two treatment conditions at day 70: Veh (exposed to the treatment vehicle dimethyl sulfoxide (DMSO) for ten days starting from day 60) and Chr (exposed to the GC dexamethasone for ten days starting from day 60). We derived the four additional treatment conditions collected at day 90 from the day 70 conditions with sustained culturing in regular media conditions for a further 20 days (wash-out period). The two conditions derived from the Veh condition were Veh-Veh and Veh-Acu, with an additional 12-hour acute GC exposure applied. Analogously, the two day-90 conditions derived from the Chr condition were Chr-Veh and Chr-Acu, with an additional 12-hour acute GC exposure applied. Figure created with BioRender.com.

We generated scRNA-seq data from all conditions. We combined these data to determine the cell type composition in organoids derived from the two iPSC lines. Basing our analysis on known marker genes of cell types in the developing human brain, we were able to identify and label eight out of the nine cell clusters in each of the lines (Fig. 2a; Supplementary Fig. 1a): Radial Glia (RG) (GPM6B, SOX2), neural progenitor cells expressing cell cycle markers (Cycling) (TOP2A, MKI67), Intermediate Progenitors (IP) (EOMES, PAX6), Excitatory Neurons (Ex. Neurons) (SLC17A6, STMN2), Inhibitory Neurons (Inh. Neurons) (GAD1, GAD2, SLC32A1, STMN2), a population of unspecified neurons expressing the G-protein regulator gene RGS5 (RGS5 Neurons) (RGS5, LINC00682, STMN2), immature Choroid Plexus cells (Imm-ChP) (RSPO3, TPBG) as well as more mature cells of the Choroid Plexus (ChP) (TTR, HTR2C, CLIC6).

**Fig. 2.**
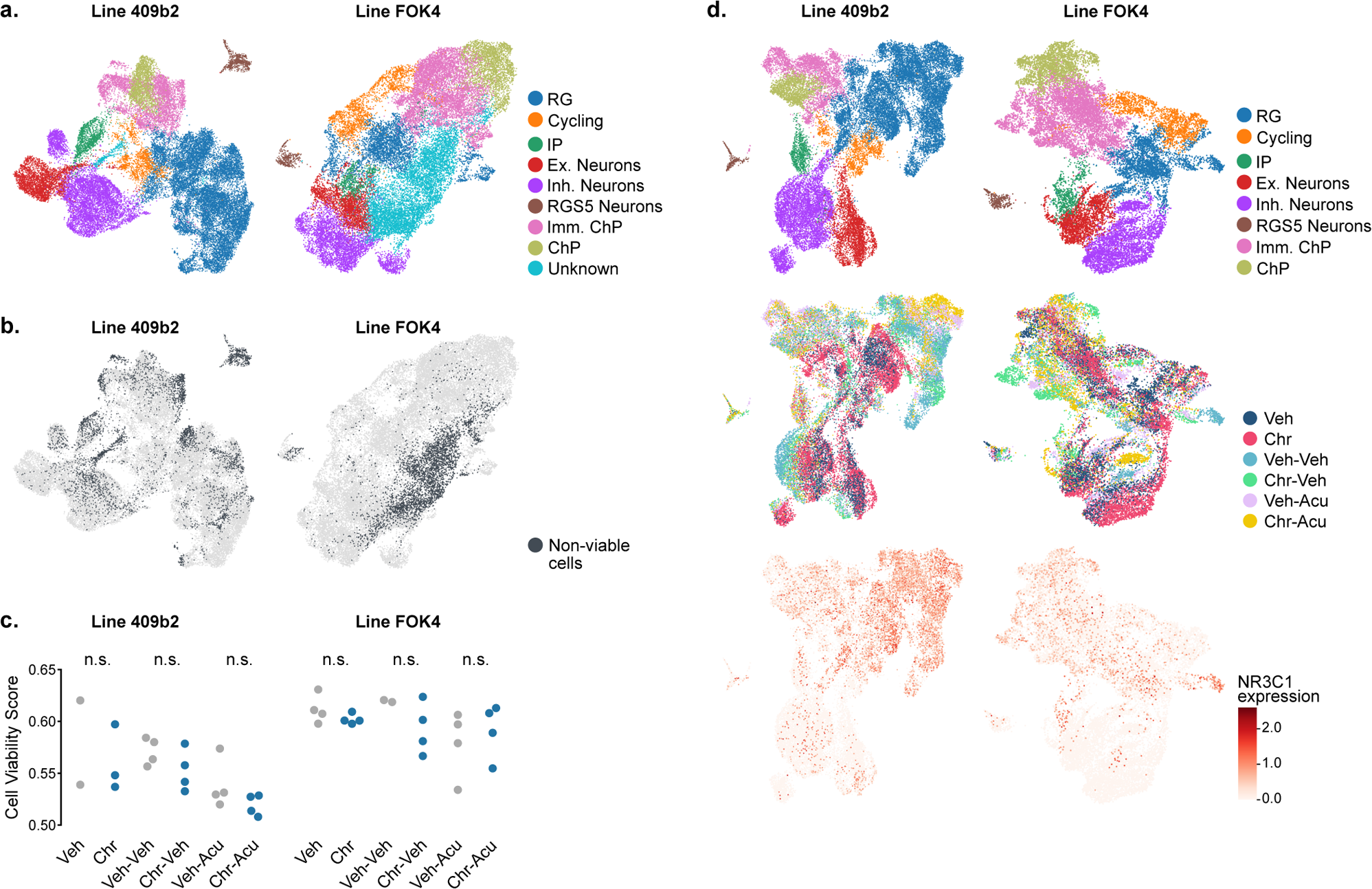
Chronic glucocorticoid exposure in neural organoids does not induce significant metabolic stress in cells. **a.** Cell types on UMAP embedding for Line 409b2 and Line FOK4. Each cell type was identified in both datasets. **b.** Localization of non-viable cells (charcoal) in the UMAP embedding for Line 409b2 and Line FOK4 organoids. The “Unknown” cluster is primarily made up of non-viable cells (74 % in Line 409b2 and 41 % in Line FOK4), suggesting these cells and the entire cluster should be removed. **c.** Swarm plots showing no significant difference in mean viability score between control (gray) and treated (blue) samples following non-viable cell removal. Each dot represents a sample from the indicated treatment condition. d. (top) UMAP embedding colored by cell type for Line 409b2 and Line FOK4 following non-viable cell removal. (middle) UMAP embedding colored by treatment conditions. (bottom) NR3C1 (GC receptor) gene expression.

We also identified a cluster of cells in the datasets from both lines, which did not express any clear combination of known marker genes. We identified many of these cells as metabolically challenged by scoring previously suggested pathways specific for non-viable cells in organoids ^29^ (Fig. 2b). We identified 12 % of all cells as non-viable in both datasets (4304 cells in Line 409b2 data and 3559 cells in Line FOK4 data). In both cell lines, we found the highest fraction of non-viable cells in the Unknown cluster (74 % of cells in Line 409b2 and 41 % in Line FOK4), followed by the RGS5 Neurons cluster (54 % of cells in Line 409b2 and 15 % in Line FOK4). Line 409b2 also had a substantial fraction of IP and Imm. ChP cells identified as non-viable cells (26 % and 20 %, respectively). For all other cell types, only small fractions of around 10 % or less were identified (Supplementary Fig. 1b). We removed all cells identified as non-viable from the datasets. We also removed the Unknown cluster in its entirety, as it was predominantly composed of non-viable cells and, beyond that, had no clear expression of marker genes. We compared viability scores between GC-exposed and control samples for the different conditions and found no significant difference after removing the non-viable cells (Fig. 2c). This finding supports that differential gene expression analyses between these conditions are likely not confounded by differences in cell viability after filtering. After removing all non-viable cells from the datasets, more evident differentiation trajectories emerged in low-dimensional space (Fig. 2d, top).

Consistent with our previous study ^22^, GC treatment was not a major source of variation during cell cluster definition, where cells were separated mainly by their cell identity in the low-dimensional data representation (Fig. 2d, middle). Furthermore, the expression of the GC receptor gene NR3C1 was uniform across most cell types but less pronounced toward the mature end of the neuronal lineage, as described previously ^22^ (Fig. 2d, bottom). To characterize the regional identity of inhibitory and excitatory neurons in our dataset, we projected them onto our recently published Human Brain Organoid Cell Atlas (HNOCA) ^30^. Our neuron subtypes projected to the non-telencephalic inhibitory and excitatory neuronal clusters in HNOCA, while our RG cells projected primarily to non-telencephalic neural progenitor cells. Organoids derived from Line 409b2 additionally contained early cells of the glial lineage, whereas organoids derived from Line FOK4 contained a more pronounced lineage of the choroid plexus. Within the neuronal lineage, cells from both lines mapped to similar cell types in HNOCA (Supplementary Fig. 1c), suggesting that organoids from both cell lines contained matching neuronal cells.

### Transcriptional response following chronic glucocorticoid treatment in organoids includes key neurodevelopmental genes

Next, we sought to identify the transcriptomic and gene-regulatory responses to our different GC treatment regimens. We harmonized results across the two iPSC donor backgrounds to ensure robustness in our identified differentially expressed (DE) genes. Specifically, we only deemed a gene as DE if it was significantly regulated at a false discovery rate smaller than 0.1 with agreeing direction of expression fold-change in organoids derived from both cell lines (Supplementary Fig. 2a). This approach reduced the number of DE genes while increasing the robustness of downstream analyses.

We first focused on DE genes in 70-day-old organoids, individually comparing Veh and Chr conditions for each identified cell type (Fig. 3a,b; Supplementary Table 1). We identified 803 consensus DE genes through this approach, with the most DE genes (n = 462) emerging in the RG cell type and the least in ChP cells with only three DE genes (Fig. 3c). Some cell types, specifically IP and RGS5+ Neurons, did not yield significant DE genes, likely due to power issues caused by smaller cell numbers in these groups or perhaps less convergence of cells within this identified cell type across the temporal differentiation timeline.

**Fig. 3.**
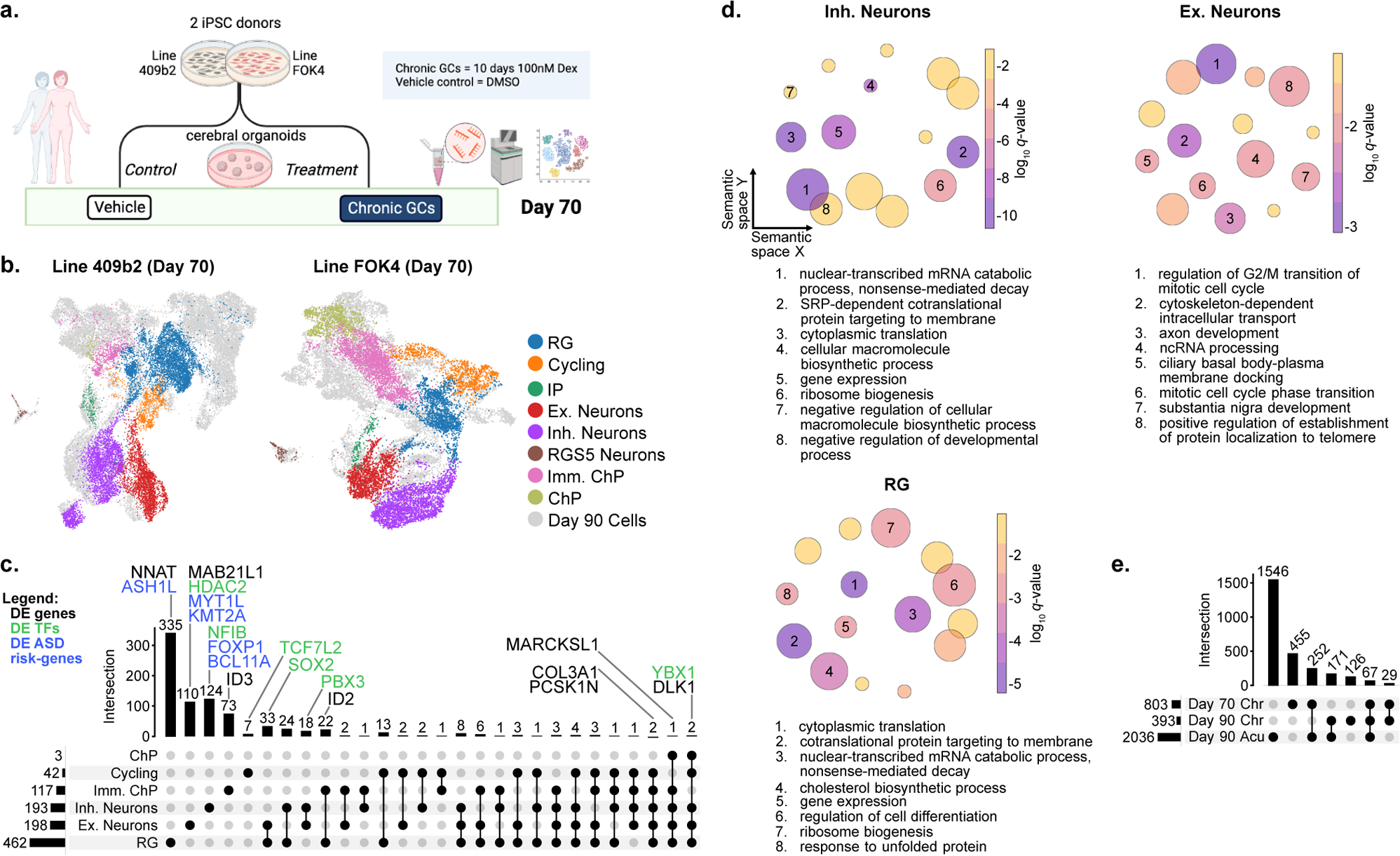
Transcriptional response following chronic glucocorticoid treatment in organoids includes key neurodevelopmental genes. **a.** Overview of the experimental design for 70-day-old organoids. Created with BioRender.com. **b.** UMAP embedding for D70 data of Line 409b2 and Line FOK4 at D70 colored by cell type. Cells from D90 samples are shown in gray. All identified cell types are present at this earlier stage. **c.** Upset plot showing consensus DE results per cell type and the number of unique and shared consensus DE genes. Selected genes are highlighted, autism risk genes are shown in blue, and further TFs are shown in green. **d.** Grouped semantic space representation of the GO-BP enrichment results for the three cell types with the most detected DE genes. The size of the circles corresponds to the number of terms in the cluster; their color corresponds to the log10(q-value) of the representative term for each cluster. The integers within the circles enumerate the eight most significant clusters, and their representative term is written out in the legend below each plot. **e.** Upset plot showing DE results aggregated across all cell types for the effect of chronic GC exposure directly following the treatment (Day 70 Chr), the effect of chronic GC exposure after a 20-day wash-out period (Day 90 Chr), and the effect of acute GC exposure in 90-day old organoids (Day 90 Acu).

Among the top consensus DE genes by fold change were NNAT, a gene associated with early neurodevelopment and ion channel control ^31^ (mean log2 fold-change (log2FC) RG = 0.56); MAB21L1, associated with cerebellum development ^32^ (mean log2FC Excitatory Neurons = 0.31); NFIB, a TF known to be essential in brain development ^33^ (mean log2FC Inhibitory Neurons = −0.50); the transcriptional regulators ID3 (mean log2FC in Imm. ChP = 0.46) and ID2 (mean log2FC in Imm. ChP and RG = 0.41 and 0.52 respectively). In addition to NFIB, we identified several additional TFs closely associated with developmental processes as DE in various clusters. Examples include SOX2, HDAC2, TCF7L2, PBX3, and YBX1. We found that one of these TFs, YBX1, was DE in a total of five cell types. Additional genes that were DE in five clusters included DLK1 (a regulator of hippocampal neurogenesis ^34^), PCSK1N (associated with the neuroendocrine system ^35^), the collagen gene COL3A1 (involved in neuronal migration ^36^), and MARCKSL1 (associated with neural tube defects and regeneration ^37^) (Fig. 3c). We next performed pathway enrichment using the Gene Ontology Biological Process (GO-BP) set. This analysis identified enrichment for terms associated with neurodevelopment in the clusters with the most consensus DE genes. These included: “axonogenesis”, “negative regulation of nervous system development” (RG); “axon development”, “substantia nigra development” (Excitatory Neurons); “cranial nerve development”, “regulation of neuron differentiation” (Inhibitory Neurons) (Supplementary Table 2). However, the terms most significantly enriched did not converge on specific neuronal pathways. They were generally associated with cell-cycle regulation and intracellular transport in Excitatory Neurons and regulation of gene expression in RG and Inhibitory Neurons (Fig. 3d).

In addition to the transcription factors mentioned to be regulated above, we also found overlap with the 36 high-risk autism spectrum disorder genes related to transcriptional regulation tested for their effect on cell fate determination using the CHOOSE system ^28^. We observed that five of these were also DE across three of our clusters: RG (ASH1L), Excitatory Neurons (MYT1L, KMT2A), and Inhibitory Neurons (FOXP1, BCL11A).

When comparing the DE effect directly following chronic GC exposure with the lasting DE effect after the wash-out period at day 90 (Supplementary Table 3), we observed a reduction in the number of DE genes after the wash-out period (393 vs. 803 consensus DE genes). With 96 DE genes shared across both comparisons, 12 % of the immediate transcriptomic effects showed persistent, stringent DE after the wash-out. The non-overlapping DE genes are either specific to a more short-lived response to GC exposure or, given the relatively high number of TFs DE at day 70, have been translated to downstream transcriptional effects, explaining the 297 newly regulated genes following the wash-out. Interestingly, 61 % of the genes regulated after chronic exposure and wash-out in 90-day-old organoids overlapped with the 2036 DE genes from the 12-hour acute stimulation at day 90 (Supplementary Table 4). Furthermore, in each cell type with overlapping genes, the directionality of the DE effect was aligned for over 90 % of the shared genes. This shows that more than half of the lasting transcriptomic effects after chronic GC exposure were shared with the response to acute GC exposure in 90-day-old organoids, supporting that the lasting effects are closely related to GC activity even after the 20-day wash-out phase. The overlap with the acute effect was prominent but reduced (40 %) in the DE day 70 genes (Fig. 3e), probably due to differences in neurodevelopmental age. In fact, within each cell type in the low-dimensional embedding, cells were separated by organoid age, emphasizing their transcriptomic differences (Supplementary Fig. 2c). We found that 13 % of the genes responsive to an acute GC exposure at day 90 had a significantly different response to this exposure based on the organoid’s history of chronic treatment, again supporting a lasting effect of the chronic GC exposure (Supplementary Table 5).

### GC exposure induces priming of the inhibitory neuron lineage in neural organoids

Having observed a DE effect in genes associated with neurodevelopment directly following treatment, particularly in a subset of high-risk genes for ASD involved in priming neuronal lineage fate ^28^, we next focused on investigating the impact of exposure to GCs on neural fate decisions within our organoid system. For this, we defined three lineage endpoints in our single-cell transcriptome data: Excitatory Neurons, Inhibitory Neurons, and Choroid Plexus (Fig. 4a). As a next step, we computed the lineage probability of every cell for each lineage endpoint along a pseudotime trajectory. For all lineage trajectories, we observed a continuous increase of lineage probability along the trajectories seen in the two-dimensional embeddings originating from early (RG) cells and reaching maximum lineage probability at the respective lineage endpoint (Fig. 4b). Given our particular interest in neuronal lineage determination and because only a few genes were DE in the ChP-related cell types, we focused on the excitatory and inhibitory neuron lineage endpoints. To account for any potential bias in organoid derivation specific to our experiments, we validated the lineage probabilities in an independently published dataset containing organoid data from 6 additional iPSC donor backgrounds ^38^. For this, we reprocessed the scRNA-seq data from 70-day-old organoids published in Kanton et al. ^38^ (Fig. 4c) and analyzed it with the same lineage inference approach used in our data. We observed similar trends in lineage determination across pseudotime with clear trajectories towards inhibitory and excitatory neurons (Fig. 4d).

**Fig. 4.**
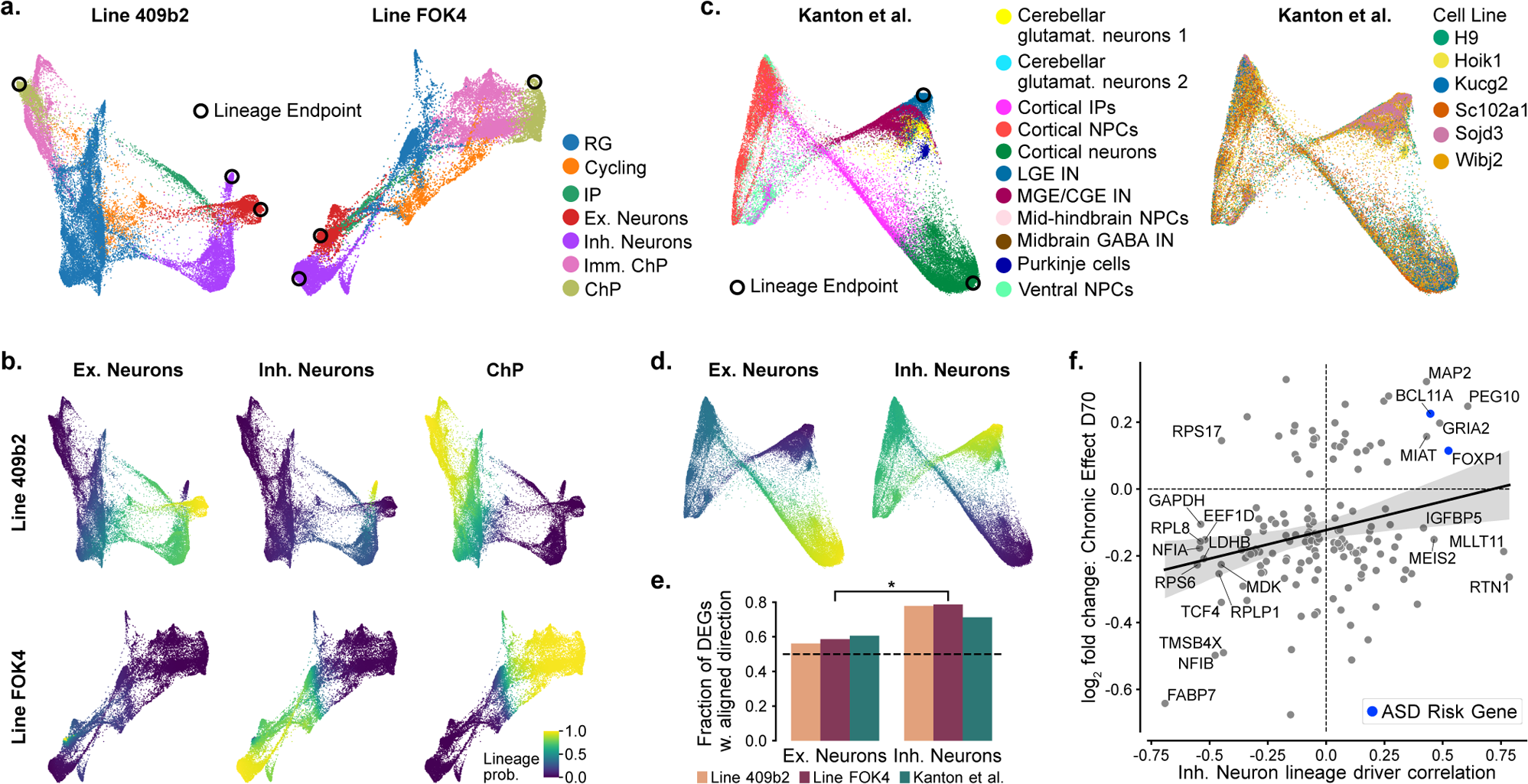
Transcription factor regulation causes priming of the inhibitory neuron lineage in neural organoids. **a.** Force-directed graph embedding of organoid data from both cell lines (without the RGS5 Neuron cluster) colored by cell type. Lineage endpoints are labeled with black circles. **b.** Computed lineage probabilities per cell for the three lineage endpoints in the two datasets. **c.** Force-directed graph layout of validation data (published 70-day-old organoid data derived from 6 additional cell lines) ^38^. Colored by cell type (left) and cell line (right). Lineage endpoints are labeled with black circles. CGE, caudal ganglionic eminence; MGE, medial ganglionic eminence; LGE, lateral ganglionic eminence; IN, interneuron; glutamat., glutamatergic; IPs, intermediate progenitors; NPCs, neural progenitor cells. **d.** Computed lineage probabilities per cell for the two neuronal lineage endpoints visualized on the force-directed graph layout of the validation data from 6 cell lines. **e.** Fraction of genes where the directionality of consensus DE effect and driver status was aligned out of all genes both significantly DE (consensus DE genes from Line 409b2 and Line FOK4 data) and in the top 500 significant driver genes (recomputed for each of the three datasets). The dashed line indicates the fraction expected by chance (0.5). **f.** Magnitude of driver gene correlation with the inhibitory neuron lineage in the validation data ^38^ vs. log2FC of consensus DE effect measured in our two cell lines. Genes with the highest lineage correlation are labeled by name, and genes associated with high risk for ASD in Li et al. ^28^ are marked in blue.

One way of defining a driver gene for a lineage is a strong correlation of the gene’s expression level with the computed lineage probability for a specific lineage endpoint ^39^. To quantify the association of the observed DE effect following GC exposure with the different lineages, we computed driver gene correlation in our two datasets and the validation dataset (8 total iPSC donors) (Supplementary Table 6). Next, we overlapped our consensus DE gene list for each neuronal cell type with the top 500 driver genes of the respective neuronal lineage from each of the three datasets. We used the resulting list of genes to compute the proportion of genes for which the directionality of the DE response aligned with the directionality of driving the lineage. For both neuronal lineages, more than half of all DE genes had an aligned direction of DE effect and expression changes along those lineages, suggesting a possible acceleration in neuronal differentiation following GC exposure. Furthermore, we found a significantly larger alignment of the DE effect with the inhibitory neuron lineage drivers in the three datasets (p=0.04) than with the drivers for the excitatory lineage. This suggested that the overall DE effect correlated to a significantly higher degree with the priming of the inhibitory than excitatory neuron lineage (Fig. 4e). Using the driver gene information from the validation dataset only, we observed a significant positive correlation between driver gene strength for the inhibitory neuron lineage and the direction of the DE effect in the inhibitory cell clusters of our two organoid datasets (r=0.25, p=0.002) (Fig. 4f). Notably, BCL11A and FOXP1, two of the genes with the strongest correlation with inhibitory neuron lineage driver genes and the largest log2FC following GC exposure, are also associated with high risk for ASD ^28^.

### GC exposure results in an increased abundance of inhibitory neurons in organoids

Next, we aimed to understand whether the observed priming of differentiation and, specifically, of the inhibitory neuron lineage translated into a measurable shift in cell type identity. We selected GAD1 as a specific marker gene of the inhibitory neuron lineage across brain regions (Fig. 5a). For both of our unguided organoid datasets, we observed an increased proportion of GAD1 positive cells at the RNA level after exposure to GCs (1.7-fold increase in Line 409b2 (5.2 % to 8.7 %); 1.2-fold increase in Line FOK4 (11.6 % to 13.8 %)) (Fig. 5b). To confirm that the increase in GAD1+ cells was not just a result of an increased number of neurons present in GC-exposed organoids, we computed the inhibitory-to-excitatory neuron ratio using GAD1 and SLC17A6 as markers for the two groups, respectively. The ratio consistently increased after GC exposure in both datasets (0.52 (Veh) to 0.86 (Chr) in Line 409b2; 2.34 (Veh) to 2.94 (Chr) in Line FOK4).

**Fig. 5.**
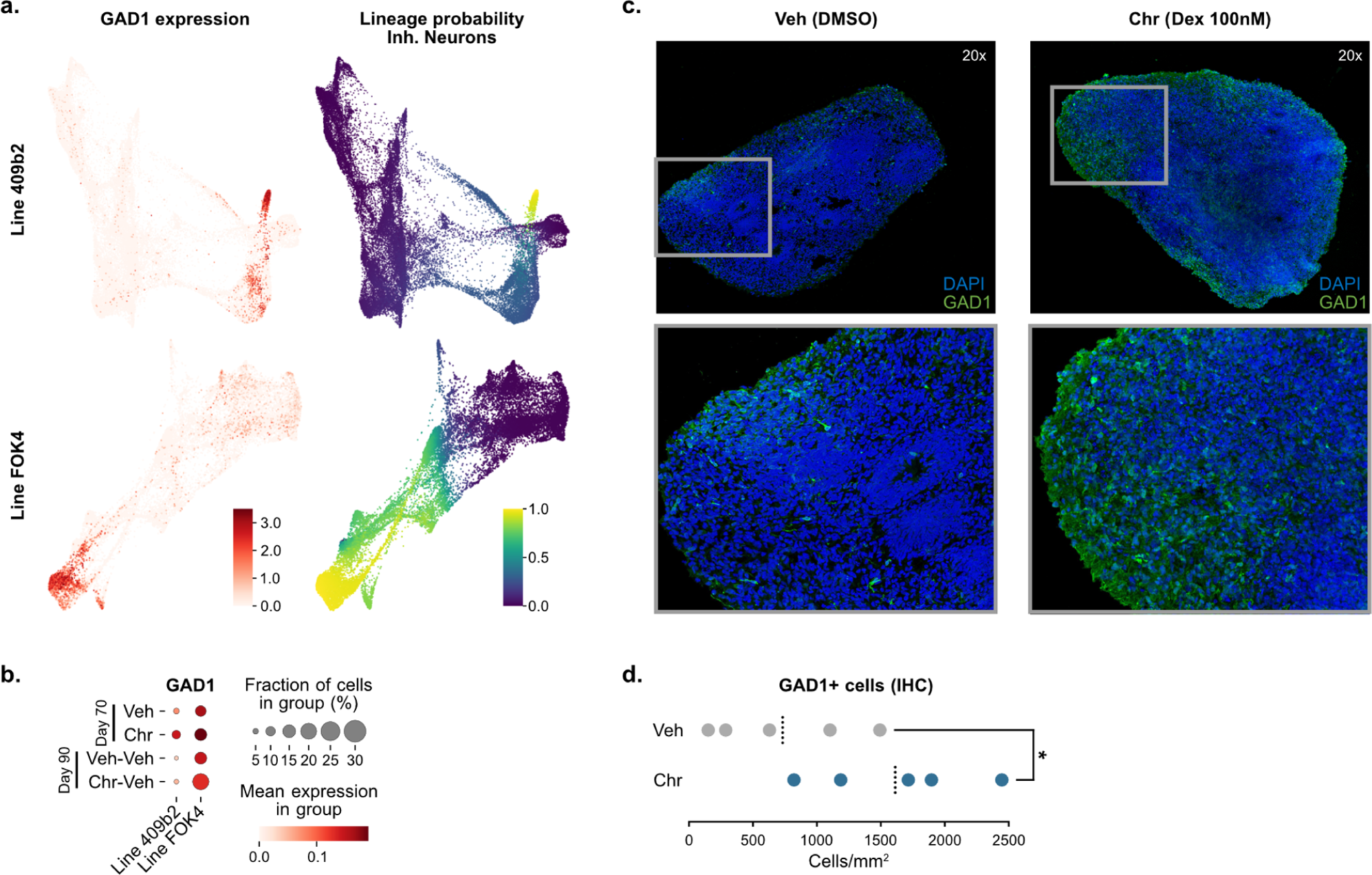
GC exposure results in an increased abundance of inhibitory neurons in organoids. **a.** Force-directed graph layout colored by expression of the inhibitory neuron marker GAD1 (left). Force-directed graph layout colored by absorption probability per cell for the inhibitory neuron lineage (right). **b.** Fraction of GAD1 positive cells and mean GAD1 expression across all cells in Veh Chr, Veh-Veh, and Chr-Veh conditions and organoids from both cell lines. **c.** Representative images of whole slice ventralized organoids at day 70 in culture, following 10 days of chronic treatment with GCs (100nM dexamethasone; Chr condition - right) and control (Veh condition - left), show an increased abundance of GAD1+ cells in the treated condition. Images were acquired at 20x magnification, showing DAPI (blue) and GAD1 (green). Lower panel: zoomed-in inserts. DMSO, dimethyl sulfoxide; Dex, dexamethasone. **d.** Cell counting quantification of GAD1+ cells across entire organoid tissue slices (n=5 per condition) and graphically represented as cells/mm^2^. Means per condition are indicated as a dotted black line. IHC, immunohistochemistry.

In order to understand whether these differences would translate to a more physiological tissue context for inhibitory neuron development, we performed a comparable exposure paradigm in regionalized ventral organoids. We obtained these stainings in a new experiment using a CRISPR/Cas9 edited version of Line 409b2 organoids expressing GFP-tagged GAD1 protein. Guiding differentiation with ventralization factors allowed for a more abundant representation of inhibitory neurons and their differentiation pathways ^40^ (Supplementary Fig. 3). Indeed, our RNA-level results were consistent with tissue-level protein expression where more GAD1 signal was apparent in 5c). Chronic exposure to GCs led to a significant 2.27-fold increase in GAD1+ cells (p=0.048, Fig. 5d, Supplementary Table 7). We replicated this finding in an independent staining experiment (fold-change=2.11; p=0.0043) (Supplementary Table 7).

Following the 20-day wash-out period, we still observed an increased number of GAD1-positive cells in 90-day-old GC-exposed organoids at the RNA level (1.2-fold increase in Line 409b2 (3.4 % to 3.9 %); 1.5-fold increase in Line FOK4 (13.5 % to 20.1 %)) (Fig. 5b), while the inhibitory-to-excitatory neuron ratio using GAD1 and SLC17A6 as markers at day 90 was not consistent across our two datasets (1.54 (Veh) to 1.15 (Chr) in Line 409b2; 5.29 (Veh) to 6.30 (Chr) in Line FOK4). In summary, we have observed priming of the inhibitory neuron lineage following GC exposure in our organoid system, resulting in more GAD1+ cells at the RNA and protein level in independent experiments, both in unguided and ventralized organoids.

### PBX3 regulation through chronic glucocorticoid exposure supports inhibitory neuron priming

We next set out to study the mechanisms underlying priming of the inhibitory neuron lineage by identifying the lineage-driving TFs with the strongest expression changes following GC exposure in our model system. We identified 15 TFs that had an aligned direction of log2FC (in inhibitory neurons following GC exposure) and driver correlation (with the inhibitory neuron lineage in the validation data) (Fig. 6a). We hypothesized that a TF that plays a central role in the lineage priming would not only itself be DE after GC exposure but would also have a significant number of its target genes DE. Across all our DE genes and all cell types, we identified 18 TFs that met these criteria: BCL6, EGR1, ID2, ID3, ID4, NEUROD1, NEUROD2, NFE2L2, NFIA, NFIB, NR2F1, NR2F2, NRG1, PBX3, SALL2, SOX2, TFAP2A, and YBX1 (Supplementary Table 8). Of these responsive TFs, the GC-upregulated hox-gene PBX3 was a positive inhibitory neuron lineage priming driver. At the same time, NFIA, NFIB, EGR1, and YBX1 were downregulated after GC exposure and negatively correlated with the inhibitory neuron lineage in the validation data. All five TFs could thus potentially contribute to the same priming effect. However, PBX3 was the only TF that consistently was among the top ten percent of driver genes for the inhibitory lineage in all three datasets (top 1.4 % - 6.5 %, depending on dataset) while not being among the top ten percent of drivers for the excitatory neuron lineage in any of the datasets (Fig. 6b, Supplementary Fig. 4a). Therefore, we focused our further analysis on PBX3 as an example of a GC-responsive TF, which could be involved in priming the inhibitory neuron lineage. We found PBX3 to be expressed across all cell types, with the highest expression levels in inhibitory neurons (Fig. 6c, Supplementary Fig. 4b). PBX3 has previously been linked to hindbrain-associated functions such as breathing, locomotion, and sensation ^41^.

**Fig. 6.**
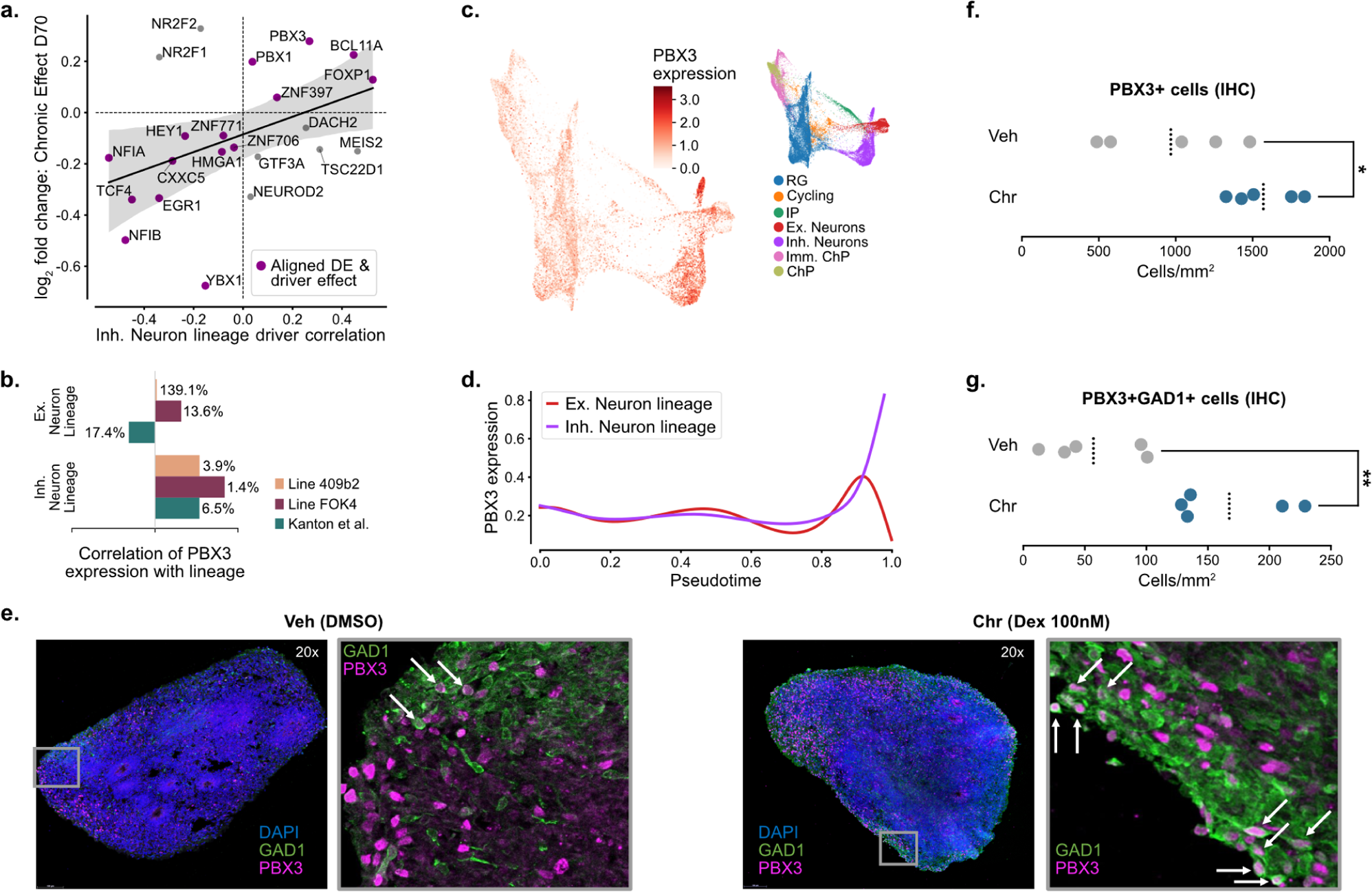
PBX3 regulation through chronic glucocorticoid exposure supports inhibitory neuron priming. **a.** Magnitude of driver gene correlation with the inhibitory neuron lineage in the validation data ^38^ vs. log2FC of GC day-70 DE effect measured in our two cell lines. Genes with aligned direction of log2FC and lineage correlation are marked in purple. **b.** Correlation of PBX3 expression with lineage probability across the excitatory and inhibitory neuronal lineages in all three datasets. The percentile of PBX3 among all significant driver genes ranked by driver strength is shown to the side of every bar. **c.** Expression of PBX3 on a force-directed graph embedding of Line 409b2 data (left) with cell type reference (right). **d.** Expression patterns of PBX3 across pseudotime for each of the two neuronal lineage endpoints in Line 409b2. **e.** Co-expression of GAD1+ (green) cells with PBX3+ (magenta) cells in 70-day-old ventralized organoids of Line 409b2 in Veh and Chr conditions. 63x magnification zoom-in images are shown to the right of the respective 20x whole slice images. Examples of double-positive cells marked by white arrows. DMSO, dimethyl sulfoxide; Dex, dexamethasone. **f.** Cell counting quantification of PBX3+ cells across entire organoid tissue slices (n=5 per condition) and graphically represented as cells/mm^2^. Means per condition are indicated as a dotted black line. IHC, immunohistochemistry. **g.** Cell counting quantification of PBX3+GAD1+ double-positive cells across entire organoid tissue slices (n=5 per condition) and graphically represented as cells/mm^2^. Means per condition are indicated as a dotted black line.

The association of PBX3 with the inhibitory lineage was supported by investigating the expression trends of PBX3 along pseudotime and towards each of the two neuronal lineage end-points, where it increased with advancing pseudotime, with the highest expression seen in inhibitory neurons. (Fig. 6d, Supplementary Fig. 4b). Exploring data from the most recent atlas of the developing human brain ^42^, we found PBX3 expression to be developmentally regulated and with overall higher expression in GABA-ergic compared to glutamatergic neurons and also higher in non-telencephalic compared to telencephalic brain regions (with highest expression in the cerebellum) (Supplementary Fig. 4c). This supports a consistent role of PBX3 in the developing fetal brain and neural organoids.

In line with a priming effect of PBX3 on inhibitory neurons, we not only observed an increased fraction of PBX3-positive cells at the transcriptional level in the Chr condition compared to the Veh condition across all cells in both cell lines (from 21 % to 26 % in Line 409b2 and from 22 % to 38 % in Line FOK4) but also an increased fraction of PBX3+GAD1+ double-positive cells (from 1.7 % to 3.9 % in Line 409b2 and from 4.5 % to 8.5 % in Line FOK4). In both cell lines, the expression of PBX3 and GAD1 was significantly positively correlated in cells expressing both genes (Line 409b2: r=0.43, p=4.1e-16; Line FOK4: r=0.36, p=2.8e-15) (Supplementary Fig. 4d).

Using immunofluorescent labeling of the PBX3 protein, we imaged the ventrally-guided organoids with GFP-tagged GAD1 at day 70 following chronic GC exposure, as previously described. We found protein-level expression patterns consistent with the scRNA-seq data, whereby PBX3 was identified preferentially (though not exclusively) in the more mature neurons that had already migrated to the outer ventricular zone (Fig. 6e). As with GAD1, we quantified PBX3+ cells across entire organoid slices and found a 1.73-fold significantly increased abundance in chronically GC-exposed organoids compared to controls (p=0.022; Fig. 6f, Supplementary Table 7). Importantly, given the apparent co-localization of the GAD1+ and PBX3+ cell populations (Fig. 6e), we also counted PBX3+GAD1+ double-positive cells and identified a 3.35-fold increase following GC exposure (p=0.0041; Fig. 6g, Supplementary Table 7). Having identified PBX3 as an example of a lineage-driving TF that responds robustly to GC treatment in our data, these results suggest a possible involvement of PBX3 in the priming of the inhibitory neuron lineage.

### Multi-modal analyses of gene regulatory networks associate PBX3 with the regulation of inhibitory neuron priming in organoids from Line 409b2

To understand the contribution of epigenetic regulation, we introduced another data modality with the same exposure paradigm. We collected scATAC-seq data for 90-day-old treated and control organoids of Line 409b2 (Veh-Veh and Chr-Veh condition). In Line 409b2, analyzed by itself, we still observed a significant positive correlation (r=0.077, p=0.046) between the DE effect in inhibitory neurons at D70 (comparing Veh to Chr condition). This correlation increased at day 90, after 20 days of wash-out (comparing Veh-Veh to Chr-Veh condition) (r=0.15, p=4.7e3), supporting a lasting effect of the treatment on the cells committed toward this lineage (Fig. 7a).

**Fig. 7.**
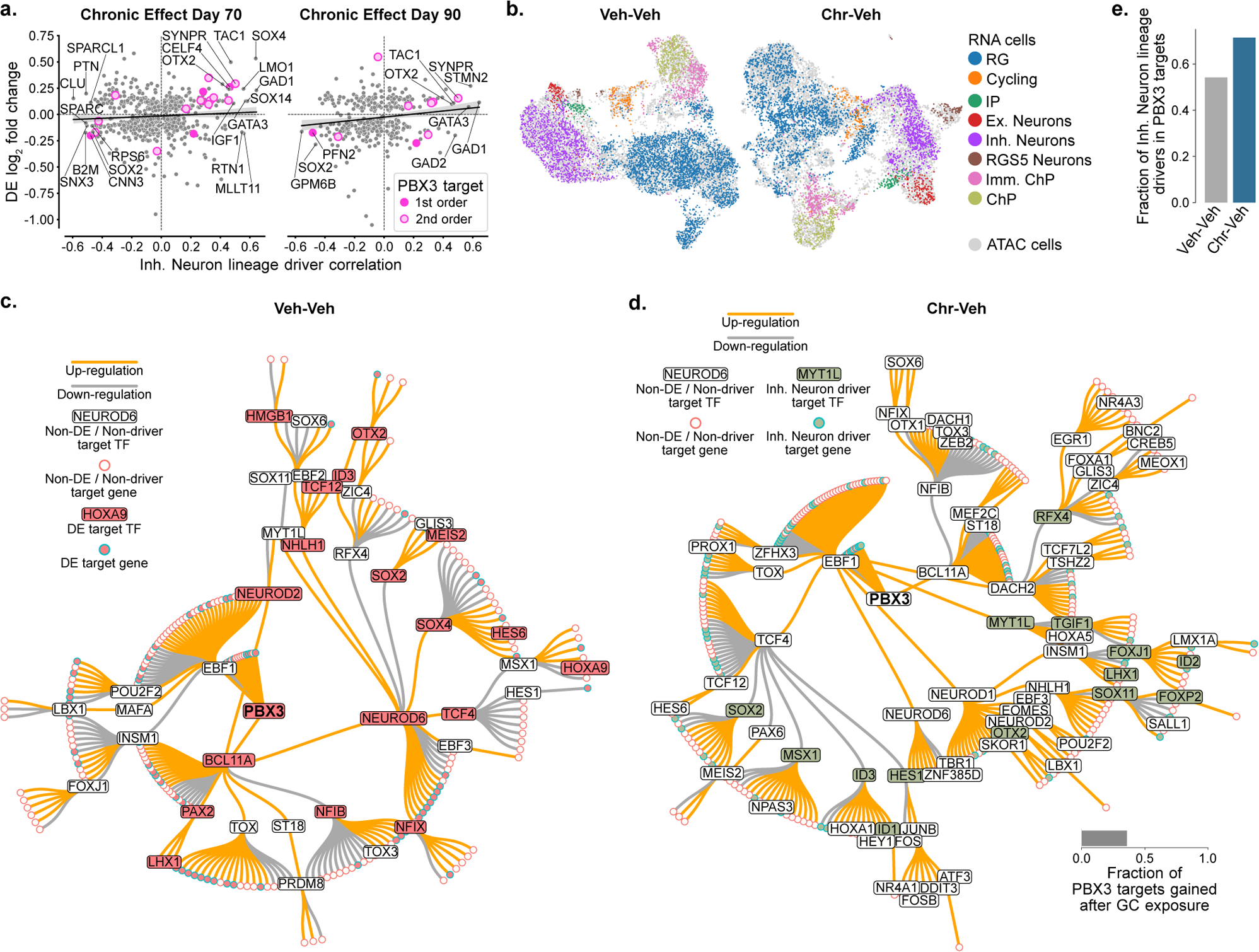
Multi-modal analyses of gene regulatory networks associate PBX3 with the regulation of inhibitory neuron priming in organoids from Line 409b2. **a.** Magnitude of driver gene correlation with the inhibitory neuron lineage vs. log2FC of Line 409b2. 1st and 2nd-order PBX3 target genes in the inferred chronic GRN are labeled in pink. Left: Directly following treatment (70 days in culture). Right: after 20 days of wash-out (90 days in culture). Genes with an absolute lineage correlation greater than 0.45 are labeled by name. **b.** UMAPS of integrated scRNAseq and scATACseq data of Line 409b2 at 90 days in culture. ScRNA-seq data is colored by cell type, and scATAC-seq data is shown in gray. Left: Vehicle organoid data. Right: GC exposed organoid data. **c.** GRN centered around PBX3 in vehicle organoids with DE genes (consensus DE genes from any of the three DE comparisons: D70 Chr, D90 Chr, D90 Acu) colored in red and TFs labeled by name. **d.** GRN centered around PBX3 in treated organoids with top 500 inhibitory neuron driver genes colored in green, TFs labeled by name. The bar chart shows the fraction of newly gained direct PBX3 downstream targets. Barplot: Fraction of inhibitory neuron drivers in direct TF downstream targets for control (Veh-Veh) and GC exposed organoids (Chr-Veh) of Line 409b2 at 90 days in culture.

Combining the scRNA-seq data with the additional scATAC-seq data enabled us to construct gene regulatory networks (GRNs) for treatment and vehicle organoids. To better understand the gene regulatory mechanisms underlying inhibitory neuron lineage priming at both vehicle and GC-exposed conditions, we integrated the single-cell genome accessibility data with the matching scRNA-seq data (Fig. 7b). We inferred multimodal GRNs using the expression of TFs and their target genes, accessibility of TF binding sites, and prior biological information such as conserved regions of the genome (Supplementary Table 9). Centering the GRN inferred in the vehicle organoids around PBX3 allowed us to visualize the baseline regulatory interactions downstream of this TF. Notably, we found that in this TF-centered GRN, 35 % of all genes were DE in at least one of the three GC treatment conditions (Veh vs. Chr, Veh-Veh vs. Chr-Veh, and Veh-Veh vs. Veh-Acu) (Fig. 7c). This again highlights the role of PBX3 in mediating the transcriptional response to GC exposure.

We computed the same PBX3-centered GRN for GC-exposed organoids and compared them to the baseline GRN. We found that 36 % of direct downstream targets in the GC-exposed condition had been gained compared to the vehicle condition, and 30 % of genes downstream of PBX3 were among the top 500 drivers of the inhibitory neuron lineage (Fig. 7d). This finding agrees with our previous observation that large fractions of the GRN are responsive to GCs and relevant for the inhibitory neuron lineage.

Comparing the association of PBX3 with the top 500 inhibitory drivers between the vehicle and GC-exposed PBX3-centered GRNs, we observed a pronounced relative increase of 32 % (from 0.54 to 0.71) in the fraction of inhibitory neuron driver genes in its direct downstream targets (Fig. 7d). In particular, some genes with a large fold-change in expression after exposure to GCs and a high correlation with the inhibitory neuron lineage were direct or second-order targets of PBX3 in the GRN of organoids exposed to GC. Examples included CELF4, a gene associated with synaptic development ^43^, depression-like behavior in mice ^44^, and ASD ^45^. Another example was SYNPR, a common inhibitory neuron marker gene ^46,47^ (Fig. 7a). Overall, these results further support a role of PBX3 in the priming of or the selection towards the inhibitory neuron lineage.

## Discussion

In this study, we investigated the effects of chronic exposure to GCs on cell type-specific gene regulation and lineage specification in neural organoids. Our experimental paradigm modeled common environmental challenges to the developing brain, specifically the prenatal administration of synthetic GCs. We observed highly cell type-specific gene expression changes directly following GC exposure that were sustained in various key molecular and cellular pathways even beyond a 20-day wash-out period. More than half of the lasting transcriptional changes after wash-out were shared with the transcriptional response to an acute GC exposure at day 90, supporting their relation to GR activation. In addition, transcripts regulated directly after chronic exposure converged on increased neuronal differentiation. We observed a positive correlation of the DE effect following GC exposure with expression changes driving the inhibitory neuron lineages, significantly more so than for excitatory neuron lineages. In line with promoting inhibitory lineage specification, in this brain model system, we showed that GC exposure leads to a higher proportion of GAD1-positive cells, a canonical marker for inhibitory neurons, that lasted following the wash-out. This suggests that GCs shift neuronal developmental trajectories from excitatory to inhibitory neuronal patterns.

We identified PBX3 as an example of an important TF for GC-induced promotion of the inhibitory neuron lineage, with both PBX3 and its downstream targets being responsive to GCs and enriched for driver genes of the inhibitory neuron lineage. In fact, GC exposure led to an increased number of cells double-positive for PBX3 and GAD1, supporting the role of this TF in GC-induced promotion of the inhibitory lineage. Interestingly, other TFs identified through this analysis to be regulated by GCs were previously identified as ASD risk genes and involved in altered inhibitory lineage specification ^28^. These included FOXP1 and BCL11A, which were up-regulated by GC and positively correlated with inhibitory neuron lineage in our dataset. This highlights the convergence of environmental and genetic risk factors on neurodevelopment, not only on specific candidate genes but also on cellular trajectories.

Previous data from our group indicated that the expression of the glucocorticoid receptor gene, NR3C1, increases in vitro until about day 40, when levels start to plateau until day 158 in organoids generated with a similar unguided protocol ^22^. Our 10-day treatment scheme (starting at day 60) and wash-out would thus fall within a window of continuously high NR3C1 expression at the whole organoid level. Previous studies in a 2D cell culture context have described that chronic exposure to GC can alter neural cell proliferation and viability ^48^. Here, neural organoids may be especially vulnerable as they have been shown to contain cells in non-physiological glycolytic or hypoxic states, likely caused by a lack of vasculature ^49,50^. Analyzing scRNA-seq data from organoids, especially in a chronic treatment context, thus requires particular care to account for metabolic profiles and altered cell states. Thus, we followed a previous study that suggested removing cells in non-physiological states to facilitate the analysis and interpretation of neural organoid scRNA-seq data ^29^. This additional quality control step allowed us to resolve the identity of an otherwise ambiguous cluster in both our datasets and improved the visualization of neuronal differentiation trajectories. It furthermore ensured that treatment and vehicle conditions had comparable cell viability scores (Fig. 2c).

Single-cell RNA-seq and scATAC-seq datasets were generated from experiments using an unguided organoid differentiation protocol, which enables the generation of a large variety of neural cell types in vitro ^51^. Regionalized organoid differentiation protocols, on the other hand, usually generate a more limited number of cell types with more regional or functional brain specificity. Notably, the exact neuronal lineages in organoids from unguided protocols can be challenging to predict a priori and do not always represent all brain regions. Therefore, a careful evaluation of the identity of obtained cell types is required so that results can be interpreted in the proper context. Mapping our datasets to the Human Neural Organoid Cell Atlas ^30^ enabled precise and efficient identification of cell types. It revealed an overlapping cell identity for non-telencephalic excitatory and inhibitory neurons, non-telencephalic neural progenitor cells, and a more limited range of glial and choroid plexus cells as expected for the brain maturity level replicated by organoids at the chosen stage.

The abundance of non-telencephalic neurons was likely an enabling factor for identifying PBX3 as a strong GC-induced inhibitory lineage driver. In a focused analysis of PBX3 expression in the developing fetal brain ^42^, we observed that this TF is developmentally regulated, with higher expression in GABA-ergic neurons and the highest levels observed in the cerebellum (Supplementary Fig. 4c). The reported effects of GCs may thus be restricted to non-telencephalic brain regions and not necessarily extrapolate to telencephalic neurons. We did, however, also find PBX3 to be a strong and specific driver of the telencephalic inhibitory neuron lineage in the dataset by Kanton et al. ^38^. Likely due to its graded expression, with highest levels in the hindbrain, including the brainstem and cerebellum, and lowest levels in the telencephalon, only very little data on the role of PBX3 in human brain development have been reported since most prior studies in vivo and in vitro, have focused on the telencephalon. However, the hindbrain has been of increasing interest in neurodevelopmental disorders, including ASD, with several studies pointing to a potentially important role of hindbrain structures, such as the cerebellum and the brainstem, in these disorders ^52,53,54^.

Our results also support the key role of the excitatory/inhibitory balance in the pathogenesis of mental and neurodevelopmental disorders (NDD) and the fact that genetic and environmental risk factors converge on this same phenotype. Previous studies in different human cellular and genetic models of ASD ^23,24,25,26,27^ and data from CRISPR-based perturbation assays in neural organoids confirm the critical role of inhibitory neurons in the pathomechanisms of these disorders. Perturbation assays targeting ASD and other NDD candidate genes have shown that they directly impact lineage specification towards inhibitory neuronal cells ^28^ and migration of inhibitory neurons ^4^. Given our findings on the effects of GC exposure on PBX3 and inhibitory neuron lineages, it would now also be important to examine the effects of other known environmental risk factors on this phenotype.

Overall, this study highlights the complex interplay between GC exposure, TF regulation, lineage specification, and neurodevelopment. It provides a molecular and cellular link between genetic and environmental risk factors for neurodevelopmental disorders, including ASD. These results also open up new avenues of investigation. Applying chronic GC treatment at different developmental time points could elucidate whether selective lineage priming is associated with varying timing of differentiation across neuronal subtypes. Probing the downstream targets and interaction partners of these TFs would provide insights into the molecular pathways involved in inhibitory lineage priming. In-vitro model systems could be used to investigate potential rescue mechanisms following lineage divergence in the context of environmental exposures. Uncovering these mechanisms can deepen our understanding of normal brain development and shed light on the molecular cascades contributing to neurodevelopmental disorders.

## Methods

### iPSC culture

Two primary human induced pluripotent stem cell (hiPSC) lines were used in this study. The first cell line was reprogrammed using hiPSCs from skin fibroblasts (HPS0076:409b2, RIKEN BRC cell bank, female) ^55,56^ and is referred to as “Line 409b2” in this manuscript. The second cell line was reprogrammed using a plasmid-based protocol for integration-free hiPSCs from peripheral blood mononuclear cells from a female donor through the BeCOME study ^57^ and is referred to as “Line FOK4” in this manuscript. MTA approvals were obtained for the use of both hiPSC lines. hiPSCs were cultured in Matrigel-coated (1:100 diluted in DMEM-F12 (Gibco™, 31330-038), Corning Incorporated, 354277) Costar® 6-well cell culture plates (Corning Incorporated, 3516) in mTESR1 Basal Medium (STEMCELL Technologies, 85851) supplemented with 1× mTESR1 Supplement (STEM-CELL Technologies, 85852) at 37°C with 5 % CO2. Passaging was performed with Gentle Cell Dissociation Reagent (STEMCELL Technologies, 07174). RevitaCell Supplement (1:100 diluted, Gibco™, A2644501) was added for 24 hours after passaging to promote cell survival.

### Neural organoid generation

Human neural organoids were created as described by Lancaster et al. ^51^ with some modifications. Briefly, hiPSCs were dissociated in StemPro Accutase Cell Dissociation Reagent (Life Technologies, A1110501). Single cells (n=9000) were dispensed into each well of an Ultra-low attachment 96-well plate with round bottom wells (Corning Incorporated, 7007) in human embryonic stem cell medium (hESC, DMEM/F12-GlutaMAX (Gibco™, 31331-028) with 20 % Knockout Serum Replacement (Gibco™, 10828-028), 3 % FBS (Fetal Bovine Serum, Gibco™, 16141-061), 1 % non-essential amino acids (Gibco™, 11140-035), 0.1 mM 2-mercaptoethanol (Gibco™, 31350-010)) supplemented with 4 ng/ml human recombinant FGF (Fibroblast Growth Factor, Peprotech, 100-18B) and 50 µM Rock inhibitor Y27632 (Millipore, SCM075) for 4 days and in hESC medium without bFGF and Rock inhibitor for an additional 2 days to form embryoid bodies (EBs). On day 6, the medium was changed to neural induction medium (NIM, DMEM/F12 GlutaMAX supplemented with 1:100 N2 supplement (Gibco™, 15502-048), 1 % Non-essential amino acids and 1 µg/ml Heparin (Sigma, H3149)) and cultured for an additional 6 days. On day 12, the EBs were embedded in Matrigel (Corning Incorporated, 354234) drops and transferred to 10cm cell culture plates (TPP, 93100) in neural differentiation medium without vitamin-A (NDM-A, DMEM/F12GlutaMAX and Neurobasal (Gibco™, 21103-049) in a 1:1 ration, additionally supplemented with 1:100 N2 supplement 1:100 B27 without Vitamin A (Gibco™, 12587-010), 0.5 % nonessential amino acids, insulin 2.5 µg/ml (Gibco™, 19278), 1:100 Antibiotic-Antimycotic (Gibco™, 15240-062) and 50 µM 2-mercaptoethanol) for 4 days. On day 16, Organoids were transferred onto an orbital shaker in NDM+A medium (same composition as NDM-A with the addition of B27 with Vitamin A (Gibco™, 17504-044) in the place of B27 without Vitamin A) and were grown in these conditions at 37°C with 5 % CO2. NDM+A medium was changed twice per week until the organoids were collected for cryopreservation or single-cell dissociation or fixation in paraformaldehyde.

For validation of inhibitory-excitatory neural lineage effects of GC, guided ventral organoids were generated as previously described by Bagley et al. ^40^. Briefly, EBs were formed starting from iPSCs dissociated into single cells using Accutase (Sigma-Aldrich, A6964) (n= 9,000). Five days later, during the neuronal induction, to induce brain regionalization, EBs were treated individually with SAG (1:10,000) (Millipore, 566660) + IWP-2 (1:2,000) (Sigma-Aldrich, I0536) for ventral identity and with cyclopamine A (1:500) (Calbiochem, 239803) for dorsal identity. All other culture parameters were identical to the ones described above for unguided organoids.

### Generation and validation of a neuron-specific fluorescent reporter iPSC cell line

Line 409b2 hiPSCs were used to generate an eGFP+/GAD1+ heterozygous iPSC cell line. gRNA (crRNA and tracrRNA, IDT) for editing with the recombinant S.p. HiFi Cas9 Nuclease V3 protein (IDT) was selected to cut efficiently at a short distance from the ATG start codon of the GAD1 gene by using the Benchling web tool (https://benchling.com). A 1611nt donor ssODNs (IDT) for homology-directed recombination was designed to have homology arms of 222-300 nt on either side of the insert DNA, a 717 nt sequence encoding for eGFP followed by the 3’UTR and the polyA signal. Lipofection (reverse transfection) was performed using the alt-CRISPR manufacturer’s protocol (IDT) with a final concentration of 10 nM of the gRNA, ssODN donor, and Cas9. In brief, 0.75 µL RNAiMAX (Invitrogen, 13778075) and the RNP mix (gRNA, ssODN, and Cas9 protein) were separately diluted in 25 µL OPTI-MEM (Gibco, 1985-062) each and incubated at room temperature for 5 min. Both dilutions were mixed to yield 50 µL of OPTI-MEM. The lipofection mix was incubated for 20–30 min at room temperature. During incubation, cells were dissociated with Accutase (Life Technologies) for 6 min and counted. The lipofection mix, 100 µL containing 50,000 dissociated cells in mTeSR1 supplemented with RevitaCell (1:100, Gibco) and the 2 µM M3814 NHEJ inhibitor ^58^ was thoroughly mixed and placed in 1 well of a 96-well plate covered with Matrigel matrix (Corning, 35248). The media was exchanged to regular mTeSR1 media (StemCell Technologies) containing the NHEJ inhibitor after 24 h. Single cell–derived clonal cell lines were analyzed and genotyped by PCR using genomic DNA isolated with QuickExtract DNA Extraction Solution (Lucigen) and primers binding within and downstream the modified region (Primer 1) or in the HAs (Primer 2).

gRNA:

5’_GGTCGAAGACGCCATCAGCT_3’

ssODN:

5’_TGCGCACCCCTACCAGGCAGGCTCGCTGCCTTTCCTCCCTCTTGTCTCTCCAGAGCCGGATCTTCAAGGGGAGCCTCCGTGCCCCCGGCTGCTCAGTCCCTCCGGTGTGCAGGACCCCGGAAGTCCTCCCCGCACAGCTCTCGCTTCTCTTTGCAGCCTGTTTCTGCGCCGGACCAGTCGAGGACTCTGGACAGTAGAGGCCCCGGGACGACCGAGCTGATGGTGAGCAAGGGCGAGGAGCTGTTCACCGGGGTGGTGCCCATCCTGGTCGAGCTGGACGGCGACGTAAACGGCCACAAGTTCAGCGTGTCCGGCGAGGGCGAGGGCGATGCCACCTACGGCAAGCTGACCCTGAAGTTCATCTGCACCACCGGCAAGCTGCCCGTGCCCTGGCCCACCCTCGTGACCACCCTGACCTACGGCGTGCAGTGCTTCAGCCGCTACCCCGACCACATGAAGCAGCACGACTTCTTCAAGTCCGCCATGCCCGAAGGCTACGTCCAGGAGCGCACCATCTTCTTCAAGGACGACGGCAACTACAAGACCCGCGCCGAGGTGAAGTTCGAGGGCGACACCCTGGTGAACCGCATCGAGCTGAAGGGCATCGACTTCAAGGAGGACGGCAACATCCTGGGGCACAAGCTGGAGTACAACTACAACAGCCACAACGTCTATATCATGGCCGACAAGCAGAAGAACGGCATCAAGGTGAACTTCAAGATCCGCCACAACATCGAGGACGGCAGCGTGCAGCTCGCCGACCACTACCAGCAGAACACCCCCATCGGCGACGGCCCCGTGCTGCTGCCCGACAACCACTACCTGAGCACCCAGTCCGCCCTGAGCAAAGACCCCAACGAGAAGCGCGATCACATGGTCCTGCTGGAGTTCGTGACCGCCGCCGGGATCACTCTCGGCATGGACGAGCTGTACAAGTAACTAGAGCTCGCTGATCAGCCTCGACTGTGCCTTCTAGTTGCCAGCCATCTGTTGTTTGCCCCTCCCCCGTGCCTTCCTTGACCCTGGAAGGTGCCACTCCCACTGTCCTTTCCTAATAAAATGAGGAAATTGCATCGCATTGTCTGAGTAGGTGTCATTCTATTCTGGGGGGTGGGGTGGGGCAGGACAGCAAGGGGGAGGATTGGGAAGACAATAGCAGGCATGCTGGGGATGCGGTGGGCTCTATGGCTTCTGAGGCGGAAAGAACCAGCTGGGGCTCTAGGGGGTATCCCCGCGTCTTCGACCCCATCTTCGTCCGCAACCTCCTCGAACGCGGGAGCGGACCCCAATACCACTAACCTGCGCCCCACAAGTAGGTCCCGCCCCAATTTTCTATCAAATGAACTGCAGGGAAGATGGGGGCGCTGGGACGTCGGGAGGCTGAGCTGGCGGAAAGGGAAGGGGGAGCGCGGAGATAATGGAGGCTGGGAAATAAATGGGGCTCTGACCCCGTCCCTGCCAGAGGTCATTCGGCTGTCAGGGACGCTAGGTGACTCCCAGGGCACCGGAAAGCGAGGACCACGCAAGGTCCGA_3’

Left and right homology arms are indicated in italics. eGFP start and stop codons are underlined.

GFP Protein translation:

MVSKGEELFTGVVPILVELDGDVNGHKFSVSGEGEGD ATYGKLTLKFICTTGKLPVPWPTLVTTLTYGVQCFSRY PDHMKQHDFFKSAMPEGYVQERTIFFKDDGNYKTRA EVKFEGDTLVNRIELKGIDFKEDGNILGHKLEYNYNSH NVYIMADKQKNGIKVNFKIRHNIEDGSVQLADHYQQN TPIGDGPVLLPDNHYLSTQSALSKDPNEKRDHMVLLEF VTAAGITLGMDELYK*

Primer 1:

For 5’_ CACTCCCACTGTCCTTTCCTAA_3’
Rev 5’_ TCCTAGCTCTTCATTCCGCC _3’

Primer 2:

For 5’:_GCTTCTCTTTGCAGCCTGTTTC_3’
Rev: 5’_ GGGCGCAGGTTAGTGGTATT _3’

### Dexamethasone treatment

Organoids were treated with glucocorticoids by dissolving Dexamethasone (Dex) in DMSO (dimethyl sulfoxide) and subsequently in the NDM+A culture medium. To achieve the final concentration of 100nM, Dex was first diluted in DMSO in a concentration of 100µM and subsequently diluted in NDM+A culture medium to a final concentration of 100nM. Vehicle control (Veh) organoids received equal amounts of DMSO. Chronic exposures (days 60-70 in culture) were performed by replacing supplemented media every two days. Some organoids were collected at day 70 following 10 days of exposure. Other organoids from the same batch were subsequently cultured in normal unsupplemented media for a 20-day wash-out period followed by an acute exposure (day 90 in culture) with 100nM Dex or DMSO for 12 hours.

### Immunoflourescence

Organoids were fixed using 4 % paraformaldehyde for 45 minutes at 4°C, cryopreserved with 30 % sucrose, fixed in optimal cutting temperature (OCT) compound (Thermo Fisher Scientific), and stored at −20°C before cutting and preparation of 16 um cryosections on SuperFrostTM slides. For immunofluorescence, sections were postfixed using 4 % PFA for 10 mins and permeabilized with 0.3 % Triton for 5 mins. Sections were subsequently blocked with 0.1 % TWEEN, 10 % Normal Goat Serum, and 3 % BSA. Primary and secondary antibodies were diluted in a blocking solution, and fluorescent staining was visualized and analyzed using a Leica laser-scanning confocal microscope. For staining with GFP and PBX3 or PAX6 and SATB2, the slides were put through antigen retrieval before fixing with paraformaldehyde. More specifically, the slides were incubated in citric buffer (0.01 M, pH 6.0) for 1 min at 720 watts and 10 mins at 120 watts, left to cool down at room temperature for 20 minutes, and washed once with PBS. Alexa anti-chicken-488 and Alexa-anti-rabbit-647 were used as secondary antibodies. All secondary antibodies are diluted to 1ug/ml or 1:1000.

#### Antibodies

| Antigen | Dilution | Vendor | Catalog # |
| --- | --- | --- | --- |
| DAPI | 1:1000 | Sigma Aldrich | D9542 |
| GFP | 1:1000 | Aves Lab | 1020 |
| PBX3 | 1:500 | Abcam | ab52903 |

### Cell imaging and counting

For immunofluorescence, stained slides were imaged using the 20x and 63x lens in confocal mode on the MICA Microhub microscope (Leica) and the Leica Application Suite X software (version 1.4.4.26810) or using the 20x lens on the AxioScan.Z1 Slide Scanner (Zeiss). To identify cells positively stained only for PBX3 and GFP-GAD1 as well as cells double-positive for PBX3 and GFP-GAD1, cells were manually counted by two independent experimenters in two separate staining experiments using the Cell Counter Tool in ImageJ software. Counts are reported as cells/mm^2^ normalized by tissue surface area. AxioScan.Z1 Slide Scanner images are quantified as total counts per entire organoid slice (n = 5 per group), while MICA Microhub images are quantified as one representative selected tile per mosaic organoid slide (n = 10 tiles per group). Statistical analyses are reported as 2-sided unpaired T-tests.

### scRNA-seq library preparation and sequencing

Single cells were dissociated using StemPro Accutase Cell Dissociation Reagent (Life Technologies), filtered through 30uM and 20uM filters (Miltenyi Biotec) and cleaned of debris using a Percoll (Sigma, P1644) gradient. Single cells were resuspended in ice-cold Phosphate-Buffered Saline (PBS) supplemented with 0.04 % Bovine Serum Albumin and prepared for single-cell separation. Experiments were performed in a paired case-control design with 2 or 4 conditions at day 70 and day 90 respectively run in parallel. Single cells were run through the Chromium controller to form gel emulsion beads containing barcoded single cells and prepared into single-cell libraries using the Chromium Single Cell 3’ Reagent Kits v2 according to the manufacturer’s recommendations without any modifications (10x Genomics). To reach an optimal target cell number, 10,000 cells per sample were loaded onto a channel of the 10x chip. All libraries were assessed using a High Sensitivity DNA Analysis Kit for the 2100 Bioanalyzer (Agilent) and KAPA Library Quantification kit for Illumina (KAPA Biosystems). Sequencing of the 10x Genomics single-cell RNA-seq libraries was performed on an Illumina NovaSeq 6000 (Illumina, San Diego, CA) at the sequencing core facility of the Max Planck Institute for Molecular Genetics (Berlin, Germany).

### scATAC-seq library preparation and sequencing

Single cells were dissociated from whole organoids according to the scRNA protocol above. Subsequent nuclei preparation and scATACseq library generation was performed using the Chromium Single Cell ATAC Library & Gel Bead Kit (16 rxns PN-1000110) according to the manufacturer’s recommendations without any modifications (10x Genomics). Sequencing of the 10x Genomics single-cell ATAC-seq libraries was performed on an Illumina NovaSeq 6000 (Illumina, San Diego, CA) at by the sequencing core facility at the Max Planck Institute for Molecular Genetics (Berlin, Germany).

### scRNA-seq quality control

Count matrices were produced from fastq files using 10x Genomics Cell Ranger ^59^ v3.0.2 with the transcriptome hg38_ensrel94^60^. Count matrices of 90-day-old acutely treated and control organoids of Line 409b2 and Line FOK4 (Veh-Veh and Veh-Acu conditions) have previously been used in Cruceanu et al. ^22^ and are available from the Gene Expression Omnibus repository (accession: GSE189534). All data has been reprocessed and reanalyzed here.

All downstream analyses were carried out using the scverse ^61^ packages scanpy ^62^ v1.9.3 and anndata ^63^ v0.9.1 with Python v3.10.12 unless indicated otherwise. For quality control (QC), cells with less than 1200 total unique molecular identifier (UMI) counts, cells with more than 150,000 total UMI counts, cells with less than 700 genes expressed, and cells with 25 % or more mitochondrial UMI counts were removed from the dataset. Next, any genes expressed in less than 20 cells were removed.

For a second QC step, an initial clustering of the full dataset was computed using louvain clustering (louvain Python package v0.8.0, https://github.com/vtraag/louvain-igraph) ^64^ with appropriate preprocessing and a resolution of 0.5. Based on this clustering, two samples, which mainly clustered separately from all other samples, were removed (409b2-D70-Chr-V1, 409b2-D70-Veh-V2). In addition, three further samples with low numbers of expressed genes or a sequencing saturation below 35 %, as reported by 10x Genomics Cell Ranger, were removed (409b2-D70-Veh-C1, FOK4-D90-Veh-Veh-C1, FOK4-D90-Veh-Veh-C2).

For a final QC step, the data was reclustered following the same procedure as before, and marker genes were computed using the gene ranking function of scanpy with default parameters. Based on the marker gene signature, any clusters containing mostly mesenchymal cells, epithelial cells, myocytes, neuroectoderm, neural stem cells, macrophages, or fibroblasts were removed from the dataset.

### scRNA-seq preprocessing and cluster annotation

All steps described in this section were applied separately to the data derived from Line 409b2 and Line FOK4.

Normalization size factors were computed using the scran ^65^ R package v1.22.1 (R v4.1.2) with the appropriate preprocessing to obtain an initial coarse clustering of the data as required for this approach. The raw counts of each cell were then normalized by the respective size factor and log(1+x) transformed. 4000 highly variable genes (HVGs) were computed using the log-normalized counts using the “cell_ranger” flavor ^59^. Principal components ^66^, a nearest neighbor graph ^67^, a force-directed graph drawing (https://github.com/bhargavchippada/forceatlas2) ^68^, louvain clustering (https://github.com/vtraag/louvain-igraph) ^64^, and partition-based graph abstraction (PAGA) ^69^ were computed using default parameters. The layout obtained from plotting the PAGA results with a threshold of 0.05 was used as initialization to compute Uniform Manifold Approximation and Projection (UMAP) ^67^ with default parameters.

Marker genes were computed by ranking genes for each cluster, with further sub-clustering or merging of clusters performed where appropriate. Cell type identities were assigned to clusters by comparing top-ranking genes per cluster with known marker genes from developmental neurobiology. A total of eight cell types and one cluster of unknown identity were identified using this procedure.

### Removing non-viable cells from scRNA-seq datasets

All steps described in this section were applied separately to the data derived from Line 409b2 and Line FOK4.

To identify cells in non-viable metabolic states, the following Gene Ontology ^70,71^ Biological Process genesets were scored using the respective scanpy function with default parameters in every cell (as previously suggested ^29^). Negative markers of cell viability: “glycolytic process” (GO:0006096), “response to endoplasmic reticulum stress” (GO:0034976). Positive markers of cell viability: “gliogenesis” (GO:0042043), “neurogenesis” (GO:0022008), and previously reported marker genes of the choroid plexus ^72^. Each score was scaled to the range (0,1). A joint cell viability score was computed by adding all scaled positive viability scores, subtracting all scaled negative viability scores, and scaling the final score to the range (0,1). Based on the final score, the “Unknown” cell cluster was identified as mostly non-viable and therefore removed from the dataset. Additionally, cells with a final viability score of less than or equal to 0.4 were identified as nonviable cells and removed from the dataset.

Following the removal of non-viable cells, HVGs, principal components, the neighbor graph, force-directed graph drawing, PAGA (threshold 0.001 for computing the layout), and UMAP were recomputed following the same procedure described in the section above.

### Mapping scRNA-seq data to the Human Neural Organoid Cell Atlas

Query to reference mapping using scPoli ^73^ from the scArches ^74^ package v0.5.9 was used to project the scRNA-seq data acquired in this study to the Human Neural Organoid Cell Atlas (HNOCA) ^30^. HNOCA data and the scPoli integration model weights used in the original study were obtained from https://github.com/theislab/neural_organoid_atlas. The feature (gene) space of the datasets from this study was adapted to the feature space used in the HNOCA scPoli model, filling any missing genes with zero expression. The query model was trained for five pre-training epochs and one training epoch with unlabeled prototype training enabled. Annealing of the model hyperparameter alpha was set to 10 epochs, and the model hyperparameter eta was set to 5. Feeding this study’s data and the HNOCA data through the trained model produced a mapped latent representation, which was used as input for the neighbor graph and UMAP computation. Cell type annotations from the annot_level_2 HNOCA annotation column (the final annotation shown in Fig. 1 of the HNOCA paper ^30^) were used to contextualize the data generated in this study.

### scRNA-seq differential expression analysis

The R tool MAST ^75^ v1.20.0 (R v4.1.2) was used to compute DE genes per cell type between treatment conditions on log-normalized expression data. This analysis was carried out separately for the two source cell lines. Samples that contained less than ten cells of a given cell type were removed to compute differential expression (DE) for this cell type. Additionally, genes expressed in less than five percent of cells of a given cell type were also removed to compute DE for this cell type. For each cell type and source cell line, a hurdle model was fit according to the following formula: *∼ ngeneson* + *treatment*_*acute* + *treatment*_*chronic* + *treatment*_*chronic treatment*_*acute* where ngeneson corresponds to the number of expressed genes in the sample, treatment_chronic corresponds to the 10-day treatment applied between day 60 and day 70 (Veh or Chr), and treatment_acute corresponds to the 12-hour treatment applied at day 90 (Veh, Acu or None for samples collected at 70 days in culture). A likelihood-ratio test was applied to test for DE at D70 (chronic effect) and D90 (chronic and acute effect).

Only genes with a false-discovery rate corrected p-value of less than 0.1 in both source cell lines and agreeing direction of DE fold-change were deemed DE for a given cell type to reduce the number of false-positive DE results. DE genes were visualized using the UpSetPlot ^76^ v0.8.0 Python package (https://github.com/jnothman/UpSetPlot).

### Functional enrichment analysis of DE genes and TF–target gene enrichment

Consensus DE genes computed as described in the previous section were used as input to the enrichment analysis using the Python implementation of Enrichr ^77,78^ via the GSEApy ^79^ package v1.0.5 with default parameters. For the annotation of biological function, the “GO_Biological_Process_2021” geneset was used as provided by Enrichr. For the transcription factor target enrichments, the “ENCODE_and_ChEA_Consensus_TFs_from_ChIP-X” geneset was used as provided by Enrichr. For greater coverage of TFs, a second database was used for this enrichment: CollectTRI ^80^, obtained via the Python implementation of decoupler ^81^ v1.4.0. Any hits with a false discovery rate of less than 0.1 were considered significantly enriched. Gene Ontology enrichment results were summarized and visualized using GO-Figure ^82^ v1.0.1 (go.obo version: releases/2021-05-01; go.obo version used to create GO relations: releases/2023-04-01; similarity_cutoff: 0.2).

### Processing of public neural organoid scRNA-seq data

Count data, associated metadata, and gene names were downloaded from ArrayExpress (accession: E-MTAB-7552) as stated in the data availability section of the Kanton et al. publication ^38^. The dataset was subsetted to cells belonging to the study’s 70-day-old organoid cell line comparison section. Any cells from cell line 409b2 or cells without a cell type label were removed from the dataset. Genes expressed in less than 10 remaining cells were also removed from the dataset. Raw counts were normalized per cell to the median total counts per cell in the dataset, and log(1 + *x*)-transformed. HVGs and principal components were computed as with the original datasets. An integrated neighbor graph was computed using the BBKNN algorithm ^83^ (bbknn Python package v1.5.1) using cell line as a batch key and neighbors_within_batch=5 with otherwise default parameters of the scanpy external implementation. From this integrated neighborhood graph, UMAP and force-directed graph drawing were computed with default parameters. The “Cortical neurons” cluster and the “LGE interneurons” cluster were identified as the excitatory and inhibitory neuron cell types in this dataset, respectively.

### Trajectory inference and driver gene computation

For Line 409b2 and Line FOK4 data, the RGS5+ Neuron cluster was removed from the dataset for this analysis, followed by recomputation of HVGs, principal components, the neighbor graph, force-directed graph drawing, PAGA (threshold 0.001 for computing the layout) and UMAP following the same procedure as described in the sections above. All the steps described in this section were applied separately to the data derived from Line 409b2, Line FOK4, and the external validation data.

The scanpy external implementation of Palantir ^84^ was used to compute a pseudo time following the manual selection of an “early cell” within a progenitor cluster for each dataset (RG for Line 409b2 and FOK4, Cortical NPCs for the validation data). First, Palantir diffusion maps were computed with five diffusion components and otherwise default parameters. Second, a t-distributed stochastic neighborhood embedding (tSNE) ^85^ representation was computed on the first two components of the Palantir multi-scale data matrix with a perplexity of 150 and otherwise default parameters. The resulting embedding was used to compute the Palantir pseudotime, sampling 500 waypoints and otherwise default parameters.

CellRank ^39,86^ was used to compute lineage probabilities based on the Palantir pseudotime. CellRank was installed from the GitHub main branch (https://github.com/theislab/cellrank) at commit c3ced63 (earliest stable version including this commit: v2.0.1). The CellRank pseudotime kernel was initiated with the Palantir pseudotime, and a transition matrix was computed. This, in turn, was used to initiate the GPCCA ^87^ estimator, which allowed the computation of macrostates from the transition matrix. Inhibitory and excitatory neuron trajectory endpoints (plus an additional ChP endpoint in Line 409b2 and FOK4 data) were manually selected from the computed macrostates and, in turn, used to compute the respective fate probabilities for each cell and terminal state. We further used CellRank to compute lineage drivers for each terminal state, correcting for false discovery rate (FDR) and discarding any drivers where the significance of the driver correlation could not be computed. A driver gene with an FDR below five percent was deemed significant in all downstream analyses. The scipy ^88^ implementation of the t-test on two related samples of scores was used to compute the significance of the difference between the alignment of consensus DE genes and driver gene directionality between the excitatory and inhibitory neuron lineages across three datasets. For visualizing gene trends, the knn-smoothing on the expression data, as implemented in the scVelo ^89^ v0.2.5 moments function, was used with 30 principal components and otherwise default parameters. Using this data, gene trends were fitted using the GAMR model ^90^ with 7 knots and plotted along Palantir pseudotime.

### Processing of public fetal human brain scRNA-seq data

The Cell-Ranger-processed count matrices from the recently published first-trimester fetal brain atlas by Braun et al. ^42^ were downloaded using the link provided by the authors (https://storage.googleapis.com/linnarsson-lab-human/human_dev_GRCh38-3.0.0.h5ad). The associated organoid age, 10x Chromium chemistry version, and neurotransmitter-transporter (NTT) annotations metadata were obtained from supplementary tables S1 and S2 of the publication: https://github.com/linnarsson-lab/developing-human-brain/files/9755355/table_S1.xlsx and https://github.com/linnarsson-lab/developing-human-brain/files/9755350/table_S2.xlsx. Any genes expressed in less than 20 cells were removed. Next, cells with less than 200 genes expressed were removed from the dataset. The total counts of each cell were normalized to 10,000 and log(1 + *x*)-transformed. Using the CellClass annotation provided by the authors, the dataset was then subset to the clusters Neuron, Neuroblast, Neuronal IPC, and Radial glia. Using the “Chemistry” annotation, the dataset was further subset to cells collected by the 10x 3’ v2 chemistry. Any cells expressing neither the GABA NTT nor any of the glutamate NTTs were removed from the dataset. Any cells expressing both the GABA NTT and any of the glutamate NTTs were also removed from the dataset. A new metadata column (“NTT_simplified”) was created, indicating whether the GABA NTT or any of the glutamate NTTs were expressed in each cell.

### scATAC-seq data preprocessing

Count matrices were produced from fastq files using 10x Genomics Cell Ranger ATAC ^91^ v2.0.0 with the reference GRCh38 (Ensembl release 94) ^60^. Unless stated otherwise, all downstream analyses were carried out using the R packages Signac ^92^ v1.9.0 and Seurat ^93^ v4.3.0 on R v4.1.2. The aggregated and filtered peak-barcode matrix from 10x Genomics Cell Ranger ATAC was loaded with Signac together with the associated fragments file and metadata. Any features detected in less than ten cells and any cells with less than 200 detected features were discarded from the dataset. Gene annotations from the EnsDb.Hsapiens.v86 v2.99.0 (https://bioconductor.org/packages/release/data/annotation/html/EnsDb.Hsapiens.v86.html) were used. Transcription start site (TSS) enrichment, nucleosome signal, and the fraction of reads in peaks statistics were computed per cell. AMULET ^94^ was installed from the GitHub main branch (https://github.com/UcarLab/AMULET) at commit 9ce413f and used for detecting and removing doublet cells from the dataset. F or QC, only cells conforming with all the following criteria were kept in the dataset: over 1,000 fragments in peak regions, less than 100,000 fragments in peak regions, TSS enrichment score greater than 2.7, TSS enrichment score smaller than 10, over 30 % reads in peaks, blacklist ration smaller than 0.66 and a nucleosome signal ratio below 10. This resulted in 7 % of cells being removed and 20616 remaining cells. TF-IDF normalization ^95^, top feature identification (min.cutoff = ‘q0’), singular value decomposition, neighbor graph computation, UMAP ^67^ computation, and clustering ^96^ were performed. Gene activities were computed and log-normalized. ChromVar ^97^ activities were computed using the BSgenome.Hsapiens.NCBI.GRCh38 (https://bioconductor.org/packages/release/data/annotation/html/BSgenome.Hsapiens.NCBI.GRCh38.html) genome and motif position frequency matrices from the JASPAR2020 database (https://bioconductor.org/packages/release/data/annotation/html/JASPAR2020.html) ^98^.

### Multimodal integration of scRNA-seq and scATAC-seq data

The integration described in this section was carried out individually for the GC-exposed and vehicle data (scRNA-seq and scATAC-seq) from 90 days-old organoids (Veh-Veh and Chr-Veh conditions) using the Python package GLUE ^99^ v0.3.2.

The scATAC-seq data was saved as an h5ad object by exporting to Python using anndata2ri v1.1 (https://github.com/theislab/anndata2ri) automatic conversion. The data was subset for the respective treatment condition and reduced to 101 dimensions using 15 latent semantic indexing iterations, as implemented in GLUE. The first dimension was discarded as it usually correlates strongly with read depth. The resulting representation was used to compute a neighbor graph as implemented in the scanpy ^62^ package (using cosine similarity as a metric), followed by UMAP ^67^ computation also using the scanpy implementation.

The raw scRNA-seq count data was subset to the respective treatment condition and processed using the scanpy package as follows, using default parameters unless stated otherwise: highly variable gene computation (n_top_genes=2000, flavor=“seurat_v3”), count normalization per cell to the median total counts per cell in the dataset, *log*(1 + *x*) transformation, scaling each feature to unit variance and zero mean, computation of 100 principle components, neighbor graph computation using the cosine similarity metric and UMAP computation.

A GLUE RNA-anchored guidance graph was computed using the scRNA-seq and scATAC-seq data, and a GLUE model was fitted using a negative-binomial probability distribution and highly variable features from both data modalities. Principal components were used as a reduced representation of the scRNA-seq data, while the latent semantic indexing embedding was used for scATAC-seq data. Data from both modalities were passed through the trained GLUE model, and the resulting concatenated representation was used to compute a combined neighbor graph and UMAP representation of the data. A bipartite matching approach ^100^, as implemented in the scim package (https://github.com/ratschlab/scim, master branch, commit 6392e65), was used (get_cost_knn_graph function with knn_k=15, null_cost_percentile=99 and capacity_method=‘uniform’) to match cells from both modalities one by one into ‘metacells’. In cases where no ATAC match was found for an RNA cell, only the RNA information was used. The GLUE latent vector of the cell was calculated as the average latent vector of the matched cells and used for joint neighbor graphs and UMAP computation for data visualization. The Python implementation of MAGIC ^101^ (https://github.com/KrishnaswamyLab/MAGIC) was used to impute gene activities on the matched dataset using k=15 neighbors, decay=1, thresh=1e-4, and four nearest neighbors for kernel bandwidth computation.

### Gene-regulatory network inference

The R tool Pando ^102^ v1.0.3 (https://github.com/quadbio/Pando) with R v4.1.2, together with Signac ^92^ v1.9.0 and Seurat ^93^ v4.3.0 for preprocessing, were used to infer gene-regulatory networks (GRNs) from the integrated multimodal data separately for the two treatment conditions (as in the integration step). The integrated metadata was loaded into a Seurat object, from where the data of the two modalities were preprocessed individually. The scATAC-seq peaks were embedded in low-dimensional space using TF-IDF normalization ^95^, top feature identification (min.cutoff = ‘q0’), and singular value decomposition. At the same time, the RNA data was log-normalized (normalization.method=“LogNormalize”, scale.factor=10000), top features were identified (selection.method=“vst”, nfeatures=4000), the data was scaled, and principle components were computed. The GRN was initiated using both data modalities and conserved regions from mammals as included in Pando (phastConsElements20Mammals.UCSC.hg38). Candidate regions were scanned for TF binding sites as provided by Pando (motif2tf data). The resulting data and initialized network were used to infer the GRN (peak_to_gene_method=’Signac’, method=’glm’) followed by gene module identification (p_thresh=0.1, nvar_thresh=2, min_genes_per_module=1, rsq_thresh=0.05). ggplot2^103^ v3.4.2 and ggraph (https://github.com/thomasp85/ggraph) v2.1.0 were used to generate TF-centered GRN visualizations.

As multimodal analyses were only carried out in Line 409b2, we applied a stricter false discovery cutoff of five percent in all DE analyses in this section to define a DE gene.

### Statistical testing

The SciPy ^88^ implementation of the t-test for the means of two independent samples was used to test for significance throughout the manuscript unless stated otherwise. Correlation coefficients were computed using the SciPy implementation of the Pearson correlation coefficient and p-value for testing non-correlation unless stated otherwise.

## Data and code availability

Processed count matrices in h5ad format (scRNA-seq and scATAC-seq) with associated metadata, as well as all analysis code is available from Zenodo. (DOI: 10.5281/zenodo.10391945; https://doi.org/10.5281/zenodo.10391945).

Raw scRNA-seq data (including filtered count matrices) are available from the Gene Expression Omnibus repository: GSE252522 (https://www.ncbi.nlm.nih.gov/geo/query/acc.cgi?acc=GSE252522): scRNA-seq Veh, Chr, Chr-Veh, and Chr-Acu conditions. GSE189534 (https://www.ncbi.nlm.nih.gov/geo/query/acc.cgi?acc=GSE189534): scRNA-seq Veh-Veh and Veh-Acu conditions; Line FOK4 = Line2; Line 409b2 = Line3; Veh-Veh = Veh, Veh-Acu = Dex. Raw scATAC-seq data (including filtered count matrices) are available from the Gene Expression Omnibus repository: GSE252523 (https://www.ncbi.nlm.nih.gov/geo/query/acc.cgi?acc=GSE252523)

## ACKNOWLEDGEMENTS

We thank Zhisong He for helpful discussions on neural cell identity in organoids and associated marker genes. We thank Marius Lange for their valuable input on lineage analysis with CellRank. We thank the sequencing core facility at the Max Planck Institute for Molecular Genetics (Berlin, Germany) for their high-throughput sequencing services. We thank the Core Facility Genomics (CF-GEN) at Helmholty Munich (Germany) for their services in processing our scRNA-seq and scATAC-seq data with Cell Ranger. This work was supported by the BMBF-funded de.NBI Cloud within the German Network for Bioinformatics Infrastructure (de.NBI) (031A532B, 031A533A, 031A533B, 031A534A, 031A535A, 031A537A, 031A537B, 031A537C, 031A537D, 031A538A) (to L.D.). L.D. acknowledges support from the Joachim Herz Foundation. C.C. acknowledges support from the Sven and Ebba-Cristina Foundation and StratNeuro. E.B.B. acknowledges support for this study from a Distinguished Investigator grant from the Brain & Behavior Research Foundation (BBRF) and funding from the Hope for Depression Research Foundation within the Depression Task Force. This preprint uses the Overleaf HenriquesLab bioRxiv template by Ricardo Henriques - CC BY 4.0 (https://www.overleaf.com/latex/templates/henriqueslab-biorxiv-template/nyprsybwffws).

## AUTHOR CONTRIBUTIONS

C.C. and A.C.K. cultured organoids, applied treatment, and generated the scRNA-seq and scATAC-seq data, with support from V.S. and M.K.; L.D. analyzed the scRNA-seq and scATAC-seq data and visualized results with input from E.B., C.C., F.J.T., S.C., A.C.K., and L.K.; M.L. generated the genetically modified GAD1-GFP iPSC line with input from S.C.; C.C., A.C.K., and C.R. generated organoids from that line and obtained immunohistochemistry stainings; C.C., I.S.D., M.G., T.S., F.B., and R.A. analyzed the immunohistochemistry data; E.B. and C.C. conceptualized the study; L.D., E.B., and C.C. wrote the manuscript with input from all co-authors. All authors read and approved the final manuscript.

## Supplementary Figures

**Supplementary Figure 1.**
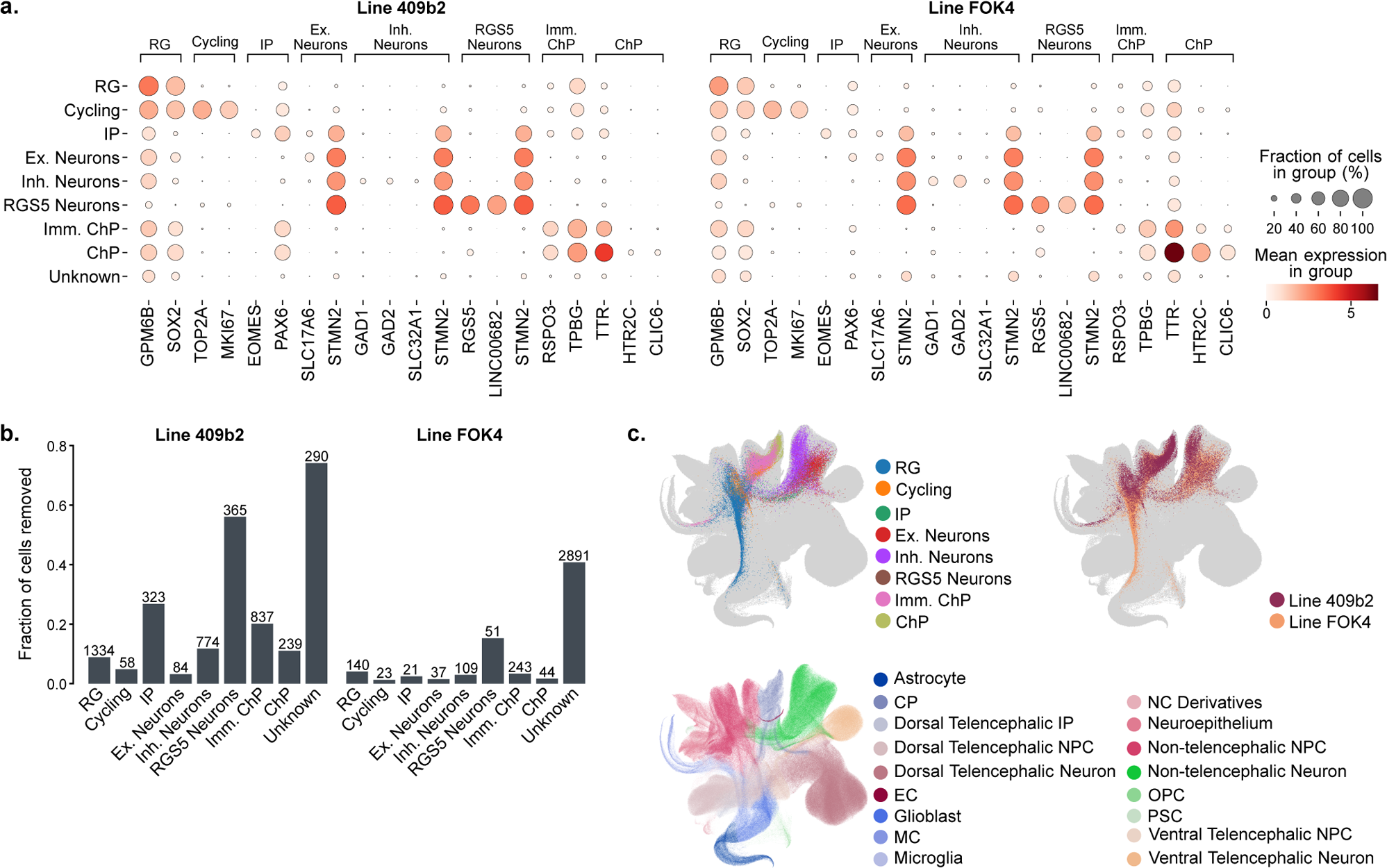
Chronic glucocorticoid exposure in neural organoids does not induce significant metabolic stress in cells. **a.** Selected marker gene expression per cell type and cell line. **b.** Fraction of non-viable cells per cell type and cell line. The absolute number of non-viable cells per cluster is displayed above each bar. The “Unknown” clusters were removed from the datasets in their entirety (394 cells in Line 409b2; 7039 cells in Line FOK4). **c.** Top: cells from this publication projected to the HNOCA ^30^. Cells are colored by their origin dataset (left) and cell types assigned in this study (right). Bottom: HNOCA cell type labels. CP, choroid plexus; NPC, neural progenitor cell; EC, endothelial cell; MC, mesenchymal cell; NC, neural crest; OPC, oligodendrocyte progenitor cell; PSC, pluripotent stem cell.

**Supplementary Figure 2.**
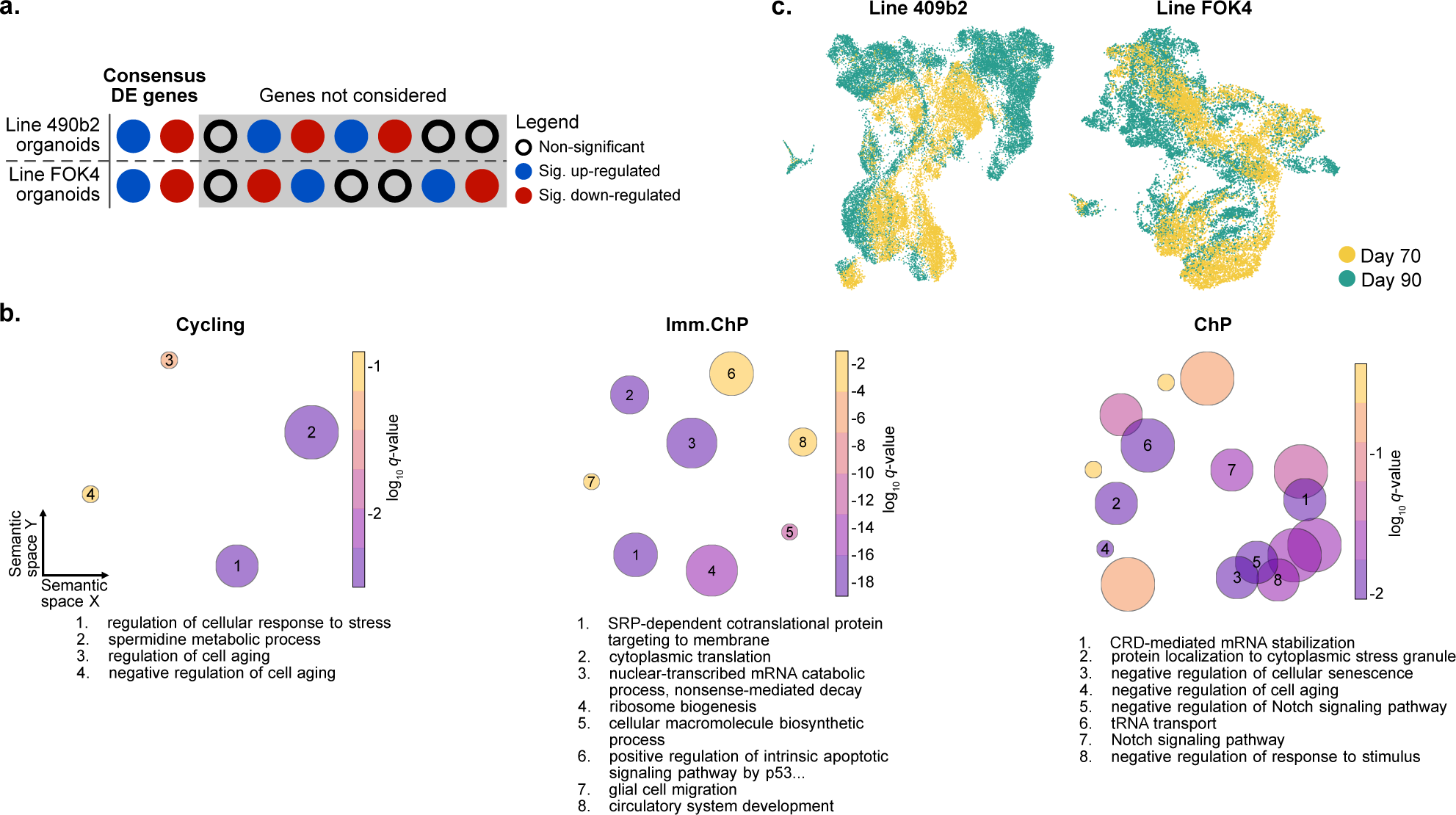
Transcriptional response following chronic glucocorticoid treatment in organoids includes key neurodevelopmental genes. **a.** Filtering scheme used to identify consensus DE genes between organoids from the two genetic backgrounds. **b.** Grouped semantic space representation of the GO-BP enrichment analysis for the three cell types with the least detected DE genes. The size of the circles corresponds to the number of terms in the cluster; their color corresponds to the log10(q-value) of the representative term for each cluster. The integers within the circles enumerate the eight most significant clusters, and their representative term is written out in the legend below each plot. **c.** UMAP embedding of Line 409b2 and Line FOK4 data colored by organoid age.

**Supplementary Figure 3.**
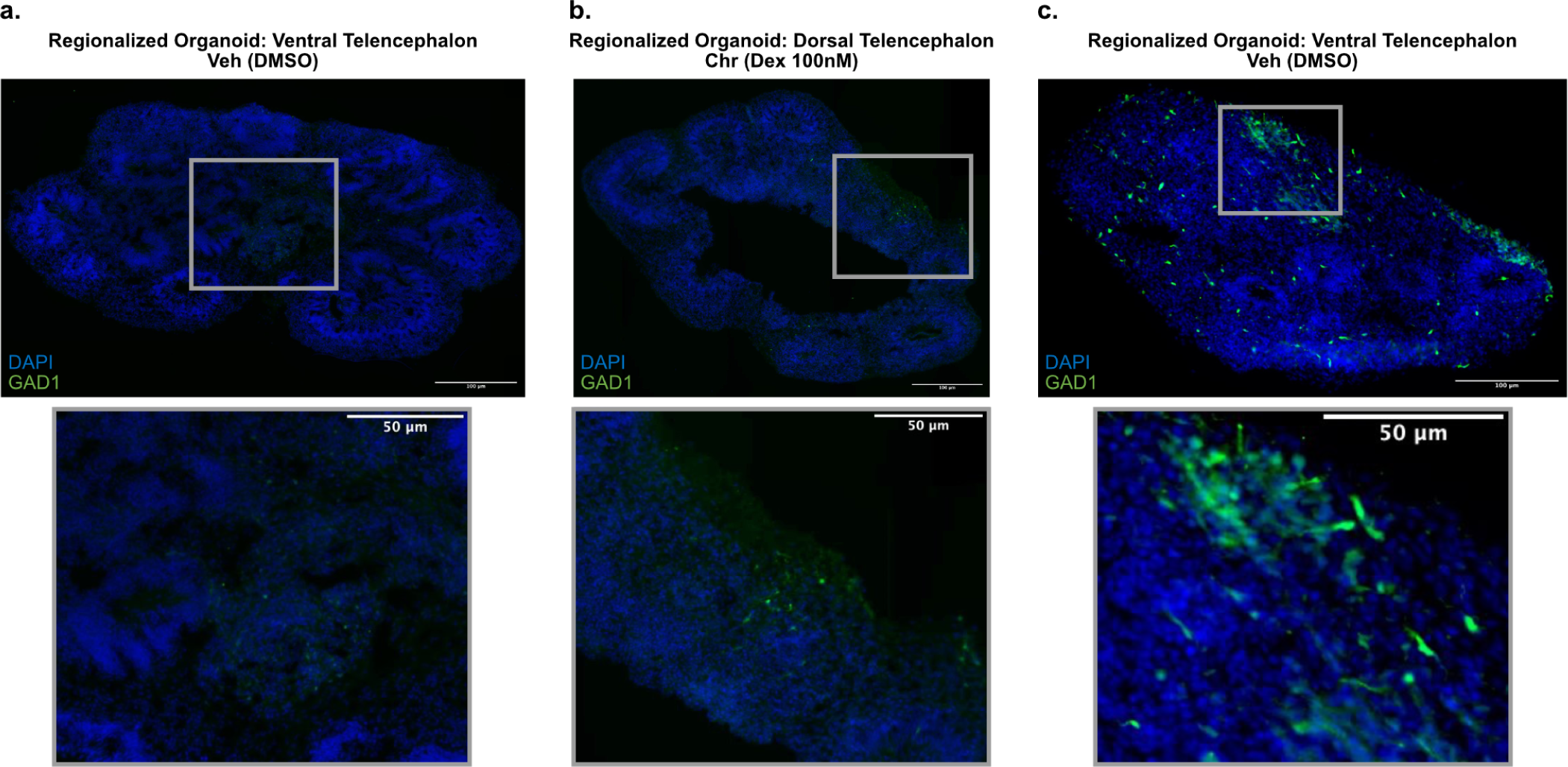
GC exposure results in an increased abundance of inhibitory neurons in organoids. **a.** Representative image of whole slice dorsalized Line 409b2 (GFP-GAD1) control organoids at day 70 in culture (Veh condition). Only very few GAD1+ cells are visible. Lower panel: zoomed-in inserts. DMSO, dimethyl sulfoxide; Dex, dexamethasone. **b.** Representative image of whole slice dorsalized Line 409b2 (GFP-GAD1) organoids at day 70 in culture, following 10 days of chronic treatment with GCs (100nM dexamethasone; Chr condition). Only very few GAD1+ cells are visible, but slightly more than in the dorsalized control organoids. **c.** Representative image of whole slice ventralized Line 409b2 (GFP-GAD1) control organoids at day 70 in culture (Veh condition). A larger number of GAD1+ cells compare to the dorsalized organoids indicates a successful ventralization.

**Supplementary Figure 4.**
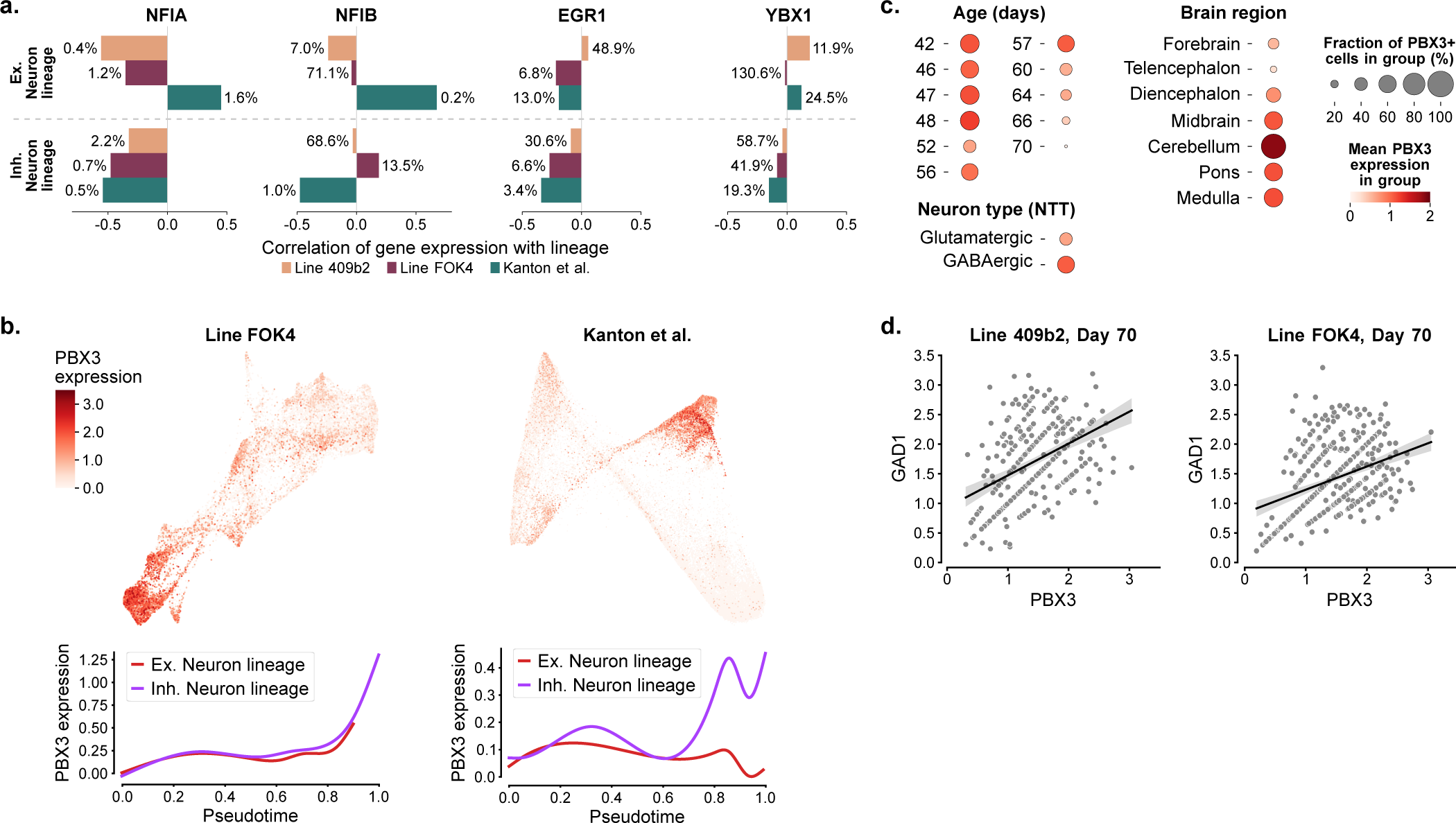
PBX3 regulation through chronic glucocorticoid exposure supports inhibitory neuron priming. **a.** Correlation of NFIA, NFIB, YBX1, and EGR1 expression with lineage probability across the excitatory and inhibitory neuronal lineages in all three datasets. The percentile of each gene among all significant driver genes ranked by driver strength is shown on the side of every bar. **b.** Expression of PBX3 in Line FOK4 and the validation data (Kanton et al.) on a force-directed graph embedding (top). Expression patterns of PBX3 across pseudotime for each of the three lineage endpoints in Line FOK4 and the validation data (Kanton et al.) (bottom). **c.** Expression of PBX3 in the fetal brain atlas ^42^ neurons and progenitors across age (in days), dissected brain region, and neurotransmitter-transporter expression (left to right). NTT, neurotransmitter transporter **d.** Expression of GAD1 and PBX3 in day-70 double-positive cells with fitted linear regression line. Left: Line 409b2. Right: Line FOK4.

## Bibliography

1. Levitt, P. & Campbell, D. B. The genetic and neurobiologic compass points toward common signaling dysfunctions in autism spectrum disorders. J. Clin. Invest. 119, 747–754 (2009).

2. Geschwind, D. H. & State, M. W. Gene hunting in autism spectrum disorder: on the path to precision medicine. Lancet Neurol. 14, 1109–1120 (2015).

3. Sullivan, P. F., Daly, M. J. & O’Donovan, M. Genetic architectures of psychiatric disorders: the emerging picture and its implications. Nat. Rev. Genet. 13, 537–551 (2012).

4. Meng, X. et al. Assembloid CRISPR screens reveal impact of disease genes in human neurodevelopment. Nature 622, 359–366 (2023).

5. Moreau, M. X., Saillour, Y., Cwetsch, A. W., Pierani, A. & Causeret, F. Single-cell transcriptomics of the early developing mouse cerebral cortex disentangle the spatial and temporal components of neuronal fate acquisition. Development 148 (2021).

6. Genovese, A. & Butler, M. G. The autism spectrum: Behavioral, psychiatric and genetic associations. Genes 14 (2023).

7. Cheroni, C., Caporale, N. & Testa, G. Autism spectrum disorder at the crossroad between genes and environment: contributions, convergences, and interactions in ASD developmental pathophysiology. Mol. Autism 11, 69 (2020).

8. Kotsiri, I. et al. Viral infections and schizophrenia: A comprehensive review. Viruses 15 (2023).

9. Davies, C. et al. Prenatal and perinatal risk and protective factors for psychosis: a systematic review and meta-analysis. Lancet Psychiatry 7, 399–410 (2020).

10. Ursini, G. et al. Placental genomic risk scores and early neurodevelopmental outcomes. Proc. Natl. Acad. Sci. U. S. A. 118 (2021).

11. Krontira, A. C., Cruceanu, C. & Binder, E. B. Glucocorticoids as mediators of adverse outcomes of prenatal stress. Trends Neurosci. 43, 394–405 (2020).

12. Painter, R. C., Roseboom, T. J. & de Rooij, S. R. Long-term effects of prenatal stress and glucocorticoid exposure. Birth Defects Res. C Embryo Today 96, 315–324 (2012).

13. Graham, A. M. et al. Maternal cortisol concentrations during pregnancy and Sex-Specific associations with neonatal amygdala connectivity and emerging internalizing behaviors. Biol. Psychiatry 85, 172–181 (2019).

14. Räikkönen, K., Gissler, M. & Kajantie, E. Maternal antenatal corticosteroid treatment and childhood mental and behavioral Disorders-Reply. JAMA 324, 1570–1571 (2020).

15. Räikkönen, K., Gissler, M. & Kajantie, E. Associations between maternal antenatal corticosteroid treatment and mental and behavioral disorders in children. JAMA 323, 1924–1933 (2020).

16. Lin, Y.-H., Lin, C.-H., Lin, M.-C., Hsu, Y.-C. & Hsu, C.-T. Antenatal corticosteroid exposure is associated with childhood mental disorders in late preterm and term infants. J. Pediatr. 253, 245–251.e2 (2023).

17. Räikkönen, K., Gissler, M., Tapiainen, T. & Kajantie, E. Associations between maternal antenatal corticosteroid treatment and psychological developmental and neurosensory disorders in children. JAMA Netw Open 5, e2228518 (2022).

18. Ninan, K., Liyanage, S. K., Murphy, K. E., Asztalos, E. V. & McDonald, S. D. Evaluation of long-term outcomes associated with preterm exposure to antenatal corticosteroids: A systematic review and meta-analysis. JAMA Pediatr. 176, e220483 (2022).

19. Tsiarli, M. A. et al. Antenatal dexamethasone exposure differentially affects distinct cortical neural progenitor cells and triggers long-term changes in murine cerebral architecture and behavior. Transl. Psychiatry 7, e1153 (2017).

20. Ohuma, E. O. et al. National, regional, and global estimates of preterm birth in 2020, with trends from 2010: a systematic analysis. Lancet 402, 1261–1271 (2023).

21. Bassil, K. et al. modeling of the neurobiological effects of glucocorticoids: A review. Neurobiol Stress 23, 100530 (2023).

22. Cruceanu, C. et al. Cell-Type-Specific impact of glucocorticoid receptor activation on the developing brain: A cerebral organoid study. Am. J. Psychiatry 179, 375–387 (2022).

23. Jourdon, A. et al. Modeling idiopathic autism in forebrain organoids reveals an imbalance of excitatory cortical neuron subtypes during early neurogenesis. Nat. Neurosci. 26, 1505–1515 (2023).

24. Mariani, J. et al. FOXG1-Dependent dysregulation of GABA/Glutamate neuron differentiation in autism spectrum disorders. Cell 162, 375–390 (2015).

25. Qian, X. et al. Sliced human cortical organoids for modeling distinct cortical layer formation. Cell Stem Cell 26, 766–781.e9 (2020).

26. Tai, D. J. C. et al. Tissue- and cell-type-specific molecular and functional signatures of 16p11.2 reciprocal genomic disorder across mouse brain and human neuronal models. Am. J. Hum. Genet. 109, 1789–1813 (2022).

27. Sawada, T. et al. Developmental excitation-inhibition imbalance underlying psychoses revealed by single-cell analyses of discordant twins-derived cerebral organoids. Mol. Psychiatry 25, 2695–2711 (2020).

28. Li, C. et al. Single-cell brain organoid screening identifies developmental defects in autism. Nature 621, 373–380 (2023).

29. Vértesy, Á. et al. Gruffi: an algorithm for computational removal of stressed cells from brain organoid transcriptomic datasets. EMBO J. 41, e111118 (2022).

30. He, Z. et al. An integrated transcriptomic cell atlas of human neural organoids. bioRxiv 2023.10.05.561097 (2023).

31. Pitale, P. M., Howse, W. & Gorbatyuk, M. Neuronatin protein in health and disease. J. Cell. Physiol. 232, 477–481 (2017).

32. Rad, A. et al. MAB21L1 loss of function causes a syndromic neurodevelopmental disorder with distinctive erebellar, cular, cranioacial and enital features (COFG syndrome). J. Med. Genet. 56, 332–339 (2019).

33. Schanze, I. et al. NFIB haploinsufficiency is associated with intellectual disability and macrocephaly. Am. J. Hum. Genet. 103, 752–768 (2018).

34. Montalbán-Loro, R. et al. dosage regulates hippocampal neurogenesis and cognition. Proc. Natl. Acad. Sci. U. S. A. 118 (2021).

35. Feng, Y., Reznik, S. E. & Fricker, L. D. ProSAAS and prohormone convertase 1 are broadly expressed during mouse development. Brain Res. Gene Expr. Patterns 1, 135–140 (2002).

36. Vandervore, L. et al. Bi-allelic variants in encoding the ligand to GPR56 are associated with cobblestone-like cortical malformation, white matter changes and cerebellar cysts. J. Med. Genet. 54, 432–440 (2017).

37. El Amri, M., Fitzgerald, U. & Schlosser, G. MARCKS and MARCKS-like proteins in development and regeneration. J. Biomed. Sci. 25, 43 (2018).

38. Kanton, S. et al. Organoid single-cell genomic atlas uncovers human-specific features of brain development. Nature 574, 418–422 (2019).

39. Lange, M. et al. CellRank for directed single-cell fate mapping. Nat. Methods 19, 159–170 (2022).

40. Bagley, J. A., Reumann, D., Bian, S., Lévi-Strauss, J. & Knoblich, J. A. Fused cerebral organoids model interactions between brain regions. Nat. Methods 14, 743–751 (2017).

41. Rottkamp, C. A., Lobur, K. J., Wladyka, C. L., Lucky, A. K. & O’Gorman, S. Pbx3 is required for normal locomotion and dorsal horn development. Dev. Biol. 314, 23–39 (2008).

42. Braun, E. et al. Comprehensive cell atlas of the first-trimester developing human brain. Science 382, eadf1226 (2023).

43. Shen, Y., Zhang, C., Xiao, K., Liu, D. & Xie, G. CELF4 regulates spine formation and depression-like behaviors of mice. Biochem. Biophys. Res. Commun. 605, 39–44 (2022).

44. Salamon, I. et al. Celf4 controls mRNA translation underlying synaptic development in the prenatal mammalian neocortex. Nat. Commun. 14, 6025 (2023).

45. Barone, R. et al. Familial 18q12.2 deletion supports the role of RNA-binding protein CELF4 in autism spectrum disorders. Am. J. Med. Genet. A 173, 1649–1655 (2017).

46. Delgado, R. N. et al. Individual human cortical progenitors can produce excitatory and inhibitory neurons. Nature 601, 397–403 (2022).

47. Blankvoort, S., Olsen, L. C. & Kentros, C. G. Single cell transcriptomic and chromatin profiles suggest layer vb is the only layer with shared excitatory cell types in the medial and lateral entorhinal cortex. Front. Neural Circuits 15, 806154 (2021).

48. Anacker, C., Zunszain, P. A., Carvalho, L. A. & Pariante, C. M. The glucocorticoid receptor: pivot of depression and of antidepressant treatment? Psychoneuroendocrinology 36, 415–425 (2011).

49. Bhaduri, A. et al. Cell stress in cortical organoids impairs molecular subtype specification. Nature 578, 142–148 (2020).

50. Giandomenico, S. L. et al. Cerebral organoids at the air-liquid interface generate diverse nerve tracts with functional output. Nat. Neurosci. 22, 669–679 (2019).

51. Lancaster, M. A. & Knoblich, J. A. Generation of cerebral organoids from human pluripotent stem cells. Nat. Protoc. 9, 2329–2340 (2014).

52. Bloomer, B. F., Morales, J. J., Bolbecker, A. R., Kim, D.-J. & Hetrick, W. P. Cerebellar structure and function in autism spectrum disorder. J Psychiatr Brain Sci 7 (2022).

53. Mapelli, L., Soda, T., D’Angelo, E. & Prestori, F. The cerebellar involvement in autism spectrum disorders: From the social brain to mouse models. Int. J. Mol. Sci. 23 (2022).

54. Bast, N., Poustka, L. & Freitag, C. M. The locus coeruleus-norepinephrine system as pacemaker of attention - a developmental mechanism of derailed attentional function in autism spectrum disorder. Eur. J. Neurosci. 47, 115–125 (2018).

55. Koyanagi-Aoi, M. et al. Differentiation-defective phenotypes revealed by large-scale analyses of human pluripotent stem cells. Proc. Natl. Acad. Sci. U. S. A. 110, 20569–20574 (2013).

56. Okita, K. et al. A more efficient method to generate integration-free human iPS cells. Nat. Methods 8, 409–412 (2011).

57. Brückl, T. M. et al. The biological classification of mental disorders (BeCOME) study: a protocol for an observational deep-phenotyping study for the identification of biological subtypes. BMC Psychiatry 20, 213 (2020).

58. Riesenberg, S. & Maricic, T. Targeting repair pathways with small molecules increases precise genome editing in pluripotent stem cells. Nat. Commun. 9, 2164 (2018).

59. Zheng, G. X. Y. et al. Massively parallel digital transcriptional profiling of single cells. Nat. Commun. 8, 14049 (2017).

60. Cunningham, F. et al. Ensembl 2019. Nucleic Acids Res. 47, D745–D751 (2019).

61. Virshup, I. et al. The scverse project provides a computational ecosystem for single-cell omics data analysis. Nat. Biotechnol. 41, 604–606 (2023).

62. Wolf, F. A., Angerer, P. & Theis, F. J. SCANPY: large-scale single-cell gene expression data analysis. Genome Biol. 19, 15 (2018).

63. Virshup, I., Rybakov, S., Theis, F. J., Angerer, P. & Wolf, F. A. anndata: Annotated data. bioRxiv 2021.12.16.473007 (2021).

64. Blondel, V. D., Guillaume, J.-L., Lambiotte, R. & Lefebvre, E. Fast unfolding of communities in large networks. J. Stat. Mech. 2008, P10008 (2008).

65. Lun, A. T. L., Bach, K. & Marioni, J. C. Pooling across cells to normalize single-cell RNA sequencing data with many zero counts. Genome Biol. 17, 75 (2016).

66. Pedregosa, F. et al. Scikit-learn: Machine learning in python. J. Mach. Learn. Res. 12, 2825–2830 (2011).

67. McInnes, L., Healy, J. & Melville, J. UMAP: Uniform manifold approximation and projection for dimension reduction. arXiv 1802.03426 (2018).

68. Jacomy, M., Venturini, T., Heymann, S. & Bastian, M. ForceAtlas2, a continuous graph layout algorithm for handy network visualization designed for the gephi software. PLoS One 9, e98679 (2014).

69. Wolf, F. A. et al. PAGA: graph abstraction reconciles clustering with trajectory inference through a topology preserving map of single cells. Genome Biol. 20, 59 (2019).

70. Ashburner, M. et al. Gene ontology: tool for the unification of biology. the gene ontology consortium. Nat. Genet. 25, 25–29 (2000).

71. Gene Ontology Consortium et al. The gene ontology knowledgebase in 2023. Genetics 224 (2023).

72. Pellegrini, L. et al. Human CNS barrier-forming organoids with cerebrospinal fluid production. Science 369 (2020).

73. De Donno, C. et al. Population-level integration of single-cell datasets enables multi-scale analysis across samples. Nat. Methods (2023).

74. Lotfollahi, M. et al. Mapping single-cell data to reference atlases by transfer learning. Nat. Biotechnol. 40, 121–130 (2022).

75. Finak, G. et al. MAST: a flexible statistical framework for assessing transcriptional changes and characterizing heterogeneity in single-cell RNA sequencing data. Genome Biol. 16, 278 (2015).

76. Lex, A., Gehlenborg, N., Strobelt, H., Vuillemot, R. & Pfister, H. UpSet: Visualization of intersecting sets. IEEE Trans. Vis. Comput. Graph. 20, 1983–1992 (2014).

77. Chen, E. Y. et al. Enrichr: interactive and collaborative HTML5 gene list enrichment analysis tool. BMC Bioinformatics 14, 128 (2013).

78. Kuleshov, M. V. et al. Enrichr: a comprehensive gene set enrichment analysis web server 2016 update. Nucleic Acids Res. 44, W90–7 (2016).

79. Fang, Z., Liu, X. & Peltz, G. GSEApy: a comprehensive package for performing gene set enrichment analysis in python. Bioinformatics 39 (2023).

80. Müller-Dott, S. et al. Expanding the coverage of regulons from high-confidence prior knowledge for accurate estimation of transcription factor activities. Nucleic Acids Res. (2023).

81. Badia-I-Mompel, P., et al. decoupler: ensemble of computational methods to infer biological activities from omics data. Bioinform Adv 2, vbac016 (2022).

82. Reijnders, M. J. M. F. & Waterhouse, R. M. Summary visualizations of gene ontology terms with GO-Figure! Front Bioinform 1, 638255 (2021).

83. Polański, K., et al. BBKNN: fast batch alignment of single cell transcriptomes. Bioinformatics 36, 964–965 (2020).

84. Setty, M. et al. Characterization of cell fate probabilities in single-cell data with palantir. Nat. Biotechnol. 37, 451–460 (2019).

85. van der Maaten, L. & Hinton, G. Visualizing data using t-SNE. J. Mach. Learn. Res. 9, 2579–2605 (2008).

86. Weiler, P., Lange, M., Klein, M., Pe’er, D. & Theis, F. Unified fate mapping in multiview single-cell data. bioRxiv 2023.07.19.549685 (2023).

87. Reuter, B., Fackeldey, K. & Weber, M. Generalized markov modeling of nonreversible molecular kinetics. J. Chem. Phys. 150, 174103 (2019).

88. Virtanen, P. et al. SciPy 1.0: fundamental algorithms for scientific computing in python. Nat. Methods 17, 261–272 (2020).

89. Bergen, V., Lange, M., Peidli, S., Wolf, F. A. & Theis, F. J. Generalizing RNA velocity to transient cell states through dynamical modeling. Nat. Biotechnol. 38, 1408–1414 (2020).

90. Wood, S. N. Fast stable restricted maximum likelihood and marginal likelihood estimation of semiparametric generalized linear models. J. R. Stat. Soc. Series B Stat. Methodol. 73, 3–36 (2011).

91. Satpathy, A. T. et al. Massively parallel single-cell chromatin landscapes of human immune cell development and intratumoral T cell exhaustion. Nat. Biotechnol. 37, 925–936 (2019).

92. Stuart, T., Srivastava, A., Madad, S., Lareau, C. A. & Satija, R. Single-cell chromatin state analysis with signac. Nat. Methods 18, 1333–1341 (2021).

93. Hao, Y. et al. Integrated analysis of multimodal single-cell data. Cell 184, 3573–3587.e29 (2021).

94. Thibodeau, A. et al. AMULET: a novel read count-based method for effective multiplet detection from single nucleus ATAC-seq data. Genome Biol. 22, 252 (2021).

95. Stuart, T. et al. Comprehensive integration of Single-Cell data. Cell 177, 1888–1902.e21 (2019).

96. Waltman, L. & Van Eck, N. J. A smart local moving algorithm for large-scale modularity-based community detection. The European physical journal B 86, 1–14 (2013).

97. Schep, A. N., Wu, B., Buenrostro, J. D. & Greenleaf, W. J. chromVAR: inferring transcription-factor-associated accessibility from single-cell epigenomic data. Nat. Methods 14, 975–978 (2017).

98. Fornes, O. et al. JASPAR 2020: update of the open-access database of transcription factor binding profiles. Nucleic Acids Res. 48, D87–D92 (2020).

99. Cao, Z.-J. & Gao, G. Multi-omics single-cell data integration and regulatory inference with graph-linked embedding. Nat. Biotechnol. 40, 1458–1466 (2022).

100. Stark, S. G. et al. SCIM: universal single-cell matching with unpaired feature sets. Bioinformatics 36, i919–i927 (2020).

101. van Dijk, D. et al. Recovering gene interactions from Single-Cell data using data diffusion. Cell 174, 716–729.e27 (2018).

102. Fleck, J. S. et al. Inferring and perturbing cell fate regulomes in human brain organoids. Nature 621, 365–372 (2023).

103. Wickham, H. ggplot2: Elegant Graphics for Data Analysis (Springer, 2016).

